# Zebrafish type I collagen mutants faithfully recapitulate human type I collagenopathies

**DOI:** 10.1101/247023

**Authors:** Charlotte Gistelinck, Ronald Y Kwon, Fransiska Malfait, Sofie Symoens, Matthew P. Harris, Katrin Henke, Shannon Fisher, Patrick Sips, Brecht Guillemyn, Jan Willem Bek, Petra Vermassen, Hanna De Saffel, MaryAnn Weis, Anne De Paepe, David R Eyre, Andy Willaert, Paul J Coucke

## Abstract

The type I collagenopathies are a group of heterogeneous connective tissue disorders, that are caused by mutations in the genes encoding type I collagen and include specific forms of Osteogenesis Imperfecta (OI) and the Ehlers-Danlos syndrome (EDS). These disorders present with a broad disease spectrum and large clinical variability of which the underlying genetic basis is still poorly understood. In this study, we systematically analyzed skeletal phenotypes in a large set of zebrafish, with diverse mutations in the genes encoding type I collagen, representing different genetic forms of human OI, and the first zebrafish model of human EDS, which harbors characteristic defects in the soft connective tissues. Furthermore, we provide insight into how zebrafish and human type I collagen are compositionally and functionally related, which is relevant in the interpretation of human type I collagen related disease models. Our studies reveal a high degree of inter-genotype variability in phenotypic expressivity that closely correlates with associated OI severity. Further, we demonstrate the potential for select mutations to give rise to variable phenotypic penetrance, mirroring the clinical variability associated with human disease pathology. Therefore, our work suggests the potential for zebrafish to aid in identifying unknown genetic modifiers and mechanisms underlying the phenotypic variability in OI and related disorders. This will improve diagnostic strategies and enable the discovery of new targetable pathways for pharmacological intervention

**SIGNIFICANCE STATEMENT:** Type I collagenopathies are a heterogenous group of connective tissue disorders, caused by genetic defects in type I collagen. Inherent to these disorders is a large clinical variability, of which the underlying molecular basis remains undefined. By systematically analyzing skeletal phenotypes in a large set of type I collagen zebrafish mutants we show that zebrafish models are able to both genocopy and phenocopy different forms of human type I collagenopathies, arguing for a similar pathogenetic basis. This study illustrates the potential of zebrafish as a tool to further dissect the molecular basis of phenotypic variability in human type I collagenopathies to improve diagnostic strategies as well as promote the discovery of new targetable pathways for pharmacological intervention of these disorders.

## INTRODUCTION

Collagens are a large family of diverse, structural extracellular matrix (ECM) proteins, among which fibril-forming molecules dominate. Type I collagen is the most abundant fibrillar collagen in tetrapods, encoded by two genes, the *COL1A1* and the *COL1A2* genes which express the α1(I) and α2(I) chain respectively. In the endoplasmic reticulum (ER), after extensive post-translational modification of proline and lysine residues, two associated proα1(I) chains and one proα2(I) chain fold into a left-handed triple helical linear molecule. The characteristic repetitive Gly-X-Y triplet sequence of the helical domain (X frequently proline, and Y frequently hydroxyproline) produces a tightly intertwined trimeric conformation (1). During the secretion of the native procollagen molecules into the ECM and assembly into fibrils, first the carboxypropeptides then the aminopropeptides are removed by propeptidases.

Genetic mutations in the type I collagen encoding genes are associated with a group of rare and heterogeneous disorders, the type I collagenopathies, which encompass specific forms of Osteogenesis Imperfecta (OI), the Ehlers-Danlos syndrome (EDS) and Caffey disease (2). Caffey disease, or infantile cortical hyperostosis, is a rare condition, caused by a specific Arg to Cys substitution (p. R836C) in the α1(I) chain of type I collagen and is characterized by an inflammatory process with swelling of soft tissues and periosteal hyperostosis of some bones (7). OI, which is a disorder characterized by a low bone mass and a high bone fragility in human patients, is predominantly caused by dominant mutations in *COL1A1* or *COL1A2* (3). According to the original classification by Sillence (4), these *COL1A* mutations underlie the *classical* subtypes of OI (type I-IV), which at biochemical level are effected by either quantitative or qualitative defects in type I collagen fibrils. OI type I, the mildest form of the disease, generally arises due to mutations that cause a premature stop codon in *COL1A1*, creating a functional null allele, so that only half of the normal amount of type I collagen is synthesized (5). The presence of one *COL1A2* null allele is not associated with a disorder, while a complete loss of *COL1A2* causes the cardiac-valvular subtype of EDS, which is associated with variable joint hypermobility and skin fragility, and early-onset severe cardiac-valvular problems, rather than bone fragility (6). In the more severe phenotypes of classical OI (OI types II, III, and IV), type I collagen exhibits a structural defect. The great majority of causative mutations in these types are single nucleotide changes, giving rise to the substitution of a Glycine residue for a bulky, polar, or charged amino acid, disrupting the highly conserved Gly-X-Y triplet sequence. Following the Sillence classification, type IV is described as being moderate, type III as severe or progressively deforming and type II as causing perinatal lethality (4). Besides the classical types of OI, mutations in non-collagenous genes have been reported to mostly cause autosomal recessive forms of OI. These genes generally encode for proteins that interact with type I collagen, acting as key players in processes such as collagen synthesis, collagen folding and post-translational modification, intra-cellular trafficking of collagen or collagen fibril cross-linking (5).

Inherent to the pathology of OI is a large clinical variability, with phenotypes ranging from nearly asymptomatic with a mild predisposition to fractures and a normal lifespan, to forms associated with multiple fractures and severe deformities already present *in utero*, that may cause perinatal lethality (7). In the classical dominant forms of OI type II-IV, some general principles have emerged for genotype-phenotype correlations (3, 8). Mutations in *COL1A1* are generally associated with a more severe phenotype than mutations in *COL1A2*. Also, the position of mutations along the α-chains can modulate the outcome. Nevertheless, numerous exceptions to these guidelines have already been demonstrated (9). Of note, extensive phenotypic variability resulting from identical mutations has been described in recent years to be common in both dominant and recessive forms of OI (10-12). This renders prediction on the clinical outcome of certain mutations in OI particularly challenging. Dissecting the underlying basis of this phenotypic variability is crucial to enable a more profound understanding of molecular mechanisms in OI, to enable the discovery of new targetable pathways for pharmacological intervention and to identify novel biomarkers for diagnostics and monitoring of therapeutic treatments.

To study human skeletal dysplasias, zebrafish (*Danio rerio*) models are increasingly being used as a valuable complementation or predecessors to the traditional murine models (13, 14). Besides their unique attributes, such as the rapid development, large offspring numbers and ease and speed in generating mutant lines, zebrafish bone mutants tend to survive into adulthood far easier than corresponding mouse models, making them also available for the study of later stages of skeletal development and maturation (13, 15). Further, due to the high conservation of developmental programs in osteogenesis between teleosts and mammals, functional gene and pathway analysis in zebrafish can yield relevant insights into human bone disease (16). Detailed reports have already documented similarities and differences between teleost and mammalian bone biology, that are relevant for modeling human bone disease (13, 14, 17, 18). One of these differences is the composition of type I collagen in zebrafish, which harbors not one but two orthologues of the human *COL1A1* gene, namely *col1a1a* and *col1a1b,* encoding the α1- and α3-chain respectively. Although some characteristics of zebrafish type I collagen have already been addressed (19), more insight into the functional implications of this additional teleost specific α-chain is needed.

In recent years, different zebrafish mutants have already been reported to accurately model certain genetic types of human OI (20-23). However, these studies where focused on detailed analysis of single mutants, modelling certain subtypes of OI, while the greatest strength of the zebrafish lies in the ease of accommodating the parallel analysis of multiple mutant models. Moreover, recent advances in microCT-based methods enable detailed and rapid skeletal phenotyping of zebrafish mutants, which often have been resistant to such methodologies (24). Advances in processing and analysis has now allowed analysis of hundreds of morphological and densitometric traits in large sets of zebrafish skeletal mutants (25). In this work, we applied systematic collagen analysis with skeletal phenomics, to characterize a large set of zebrafish with mutations in type I collagen genes as seen in patients with different forms of classical OI and EDS, to asses to which extent key features of human type I collagenopathies are recapitulated.

## MATERIAL AND METHODS

### Animals

The *col1a1a*^sa1748^, *col1a1b*^sa12931^*, col1a2*^sa17981^*, bmp1a*^sa2416^ and *plod2*^sa1768^ mutant zebrafish were generated by the zebrafish mutation project (ZMP) and together with wild-type AB fish obtained from the zebrafish international resource center (ZIRC, http://zebrafish.org) or the European Zebrafish Resource center (EZRC, http://www.ezrc.kit.edu) (26). The *col1a1a*^chi^ were previously described (21) and the *col1a1a*^med^ mutant fish were purchased from the EZRC. The *col1a1a*^dmh13^, *col1a1a*^dmh14^*, col1a1b*^dmh29^ and *col1a2*^dmh15^ mutant fish were generated in a forward genetics screen (27). Fish were housed in ZebTEC semi-closed recirculation housing systems (Techniplast, Italy) and were kept at a constant temperature (27-28°C), pH (7.5) and conductivity (500 µS) on a 14/10 light/dark cycle. Zebrafish larvae were polycultured in static tanks with rotifers for the first 5 days of feeding (days 5-9 postfertilization) according to Best *et al.* (28). Fish were fed three times a day with both dry food (Skretting and Zebrafish Management Ltd., UK) and micro-artemia (Ocean Nutrition, Belgium) (28). Embryos were collected by natural spawning, staged and raised according to Kimmel and colleagues (29), in agreement with EU Directive 2010/63/EU for animals, permit Number: ECD 17/68. All efforts were made to minimize pain and discomfort.

### µCT scanning and analysis

For µCT-based phenotyping and quantification, adult zebrafish were euthanized using 0.4% tricaine, stored frozen, and thawed immediately prior to scanning. Whole body µCT scans of 5 mutant and 5 control fish of each genotype were acquired on a Scanco Viva CT 40 (Scanco, Switzerland) using the following scan parameters: 55kV, 145µA, 200ms integration time, 500 proj/180°, and 21µm voxel size. Two fish were scanned simultaneously in each acquisition and DICOM files of individual fish were generated using Scanco software. DICOM files of individual fish were segmented in MATLAB using custom FishCuT software and data were analyzed in the R statistical environment, as previously described (25). The global test was used to assess whether the pattern across vertebrae is significantly different between a certain mutant genotype and its control group of WT siblings.

### Collagen analysis

Adult zebrafish were euthanized with 0.4% tricaine and spine fragments of 3 mutant and 3 control fish for each genotype were isolated by manual dissection, residual tissue was removed by incubation in Accumax solution (Sigma-Aldrich, USA) for 1 hour. Bone was demineralized in 0.1 M HCl at 4°C, washed, and solubilized by heat denaturation in SDS-PAGE sample buffer. The method of Laemmli was used with 6% gels for the denaturant extracts. Collagen α-chains were cut from SDS-PAGE and subjected to in-gel trypsin digestion. Electrospray MS was performed on tryptic peptides using an LTQ XL ion-trap mass spectrometer (Thermo Scientific, USA) equipped with in-line liquid chromatography using a C4 5*µ*m capillary column (300um × 150mm; Higgins Analytical RS-15M3-W045) eluted at 4 *µ*l/min. The LC mobile phase consisted of buffer A (0.1% formic acid in MilliQ water) and buffer B (0.1% formic acid in 3:1 acetonitrile:n-propanol v/v). The LC sample stream was introduced into the mass spectrometer by electrospray ionization (ESI) with a spray voltage of 4kV. Proteome Discoverer search software (Thermo Scientific, USA) was used for peptide identification using the NCBI protein database. In order to determine a relative mount of α1(1)- and α3(1)-chains a unique tryptic peptide for each α1(1) and for α3(1) was selected and the relative amount of the corresponding peak manually measured on the full scan by determining peak height. For all mutant genotypes an expected ratio of α3(I)/α1(I) was calculated based on the mean relative α3(I)/α1(I) ratio observed in control samples and based on the assumption of absence of α3(I) and α1(I) in case of knockout of respectively *col1a1b* and *col1a1a*.

### Histological analysis

For paraffin embedding, adult zebrafish were first euthanized using 0.4% tricaine. An incision was made in the abdomen of the fish from the gills to the anal pore and zebrafish were fixed overnight in modified Davidson’s Fixative. The solution was changed to 10% NBF (VWR Chemicals, USA) and incubated overnight. For decalcification, the solution was changed to citric acid (1 volume 45% formic acid, 1 volume 20% sodium citrate, 2 volumes water) and incubated for 4 hours. Subsequently the zebrafish were dehydrated overnight in 70% EtOH, followed by 90% EtOH for 2 hours and finally 100% EtOH overnight on a shaker. Samples were placed in xylene for 8 hours in a fume hood. The solution was then changed to paraffin and samples were placed at 65°C overnight. Lids were removed to allow the remaining xylene to evaporate. Samples were embedded in paraffin. Subsequently, tissues were sectioned at 8um thickness and stained with Masson’s trichrome staining (Sigma-Aldrich, USA, HT15-1KT) following guidelines outlined by the manufacturer. All sections were examined and photographed using a Leica DM 2000 Led microscope (Leica, Germany). of the corresponding peak manually measured on the full scan by determining peak height.

### Bio-mechanical testing

Soft connective tissue sections were isolated from 5 months old *col1a2*^−/−^ mutant fish (n=5) and wild type control siblings (n=5) by decapitation and dissecting away the spine from each specimen. These tissue specimens were kept in PBS for a maximum period of 8 hours. Uniaxial tensile tests were performed at room temperature using an Instron^®^ 5942 electromechanical test system (Instron, USA). The system was configured with lightweight BioPuls pneumatic side action grips and surfalloy jaw faces. Tissue specimens were tested with a 10N full-scale load cell, and were loaded onto the machine with 7mm distance between the two vertical clamps. Tissues were stretched at a constant displacement rate of 12mm/min until rupture or maximum load/extension. The load and deformation values were recorded continuously for each strain increment of 0.2%. The breaking strength was defined as the load required to break the specimen, called maximum load (30).

## RESULTS

### Zebrafish with different mutations in type I collagen show variable skeletal phenotypes

We systematically analyzed skeletal phenotypes in a large set of zebrafish models carrying different mutations in the zebrafish type I collagen encoding genes *col1a1a, col1a1b* and *col1a2* (Table 1). As a reference, we also included two knock-out mutants, *bmp1a*^*−/−*^ and *plod2*^*-/*-^, representing severe recessive forms of OI with defects in type I collagen processing and cross-linking respectively (22). Upon microCT scanning we observed a large diversity of skeletal phenotypes throughout the whole set of mutants. Severe morphological abnormalities included callus formation, bowing and kinking of the ribs, malformation of the vertebral column, and short stature (Figure 1, Supplementary Table 1), indicating compromised bone quality. Next, we performed microCT-based phenomic profiling (25) to quantify 200 different descriptors of bone morphology and mineralization in the axial skeleton of each animal. Specifically, we segmented each of 20 vertebra into three skeletal elements: the neural arch (Neur), centrum (Cent), and haemal arch/ribs (Haem), and for each skeletal element, we computed four primary parameters: Tissue Mineral Density (TMD, mgHA/cm3), Volume (Vol, µm3) and Thickness (Th, µm), and also centrum length (Cent.Le, µm). In total, we analyzed 28000 phenotypic data points derived from 140 different animals. To explore inter-genotype variability for each trait, means were calculated within each genotype (mutant and control), and normalized to the standard deviation of their respective control population (Z-score = ((mean value mutant – mean value control)/SD Control). In general, we observed a large diversity of skeletal phenotypes, throughout the whole set of mutants (Figure 2). In general, we found that phenotypic severity (as measured by the absolute value of the Z-score) tended to be highest for mutants associated with collagen processing (*plod2*, *bmp1a*) and qualitative collagen defects (e.g., *col1a1a*^*chi/+*^, *col1a2*^*dmh15/+*^, *col1a1b*^*dm29/+*^), and lower for mutants associated with quantitative collagen defects (e.g., *col1a1a*^+/−^, *col1a1b*^−/−^, *col1a2*^−/−^;). Remarkably, some mutants with qualitative defects (and collagen processing defects) exhibited a pronounced enrichment of positive Z-scores for some traits (e.g., Cent.TMD). We interpret the different degrees of phenotypic severity in mutants associated with qualitative or quantitative collagen defect, as well as the variability in direction of effect (i.e., positive or negative effect on bone mass or mineralization) across genotypes, to reflect the clinical variability observed in human type I collagenopathies.

**Table 1.** List of mutant zebrafish alleles analyzed in this study.

| Gene | Allele | Effect on protein level | Effect on homologous human protein |
| --- | --- | --- | --- |
| Type I collagen knockout alleles (quantitative defect) |  |  |  |
| *col1a1a* | *sa1748* | p.(Gly1179X) |  |
| *col1a1b* | *sa12931* | p.(Cys68X) |  |
| *col1a2* | *sa17981* | p.(Ala154Cysfs*23) |  |
| Type I collagen amino-acid substitutions (qualitative defect) |  |  |  |
| *col1a1a* | *chihuahua (chi)* | p.(Gly736Asp) | α1(I) p.(Gly574Asp) |
| *col1a1a* | *microwaved (med)* | p.(Glu888Lys) | α1(I) p.(Glu726Arg) |
| *col1a1a* | *dmh13* | p.(Gly1093Arg) | α1(I) p.(Gly931Arg) |
| *col1a1a* | *dmh14* | p.(Gly1144Glu) | α1(I) p.(Gly981Glu) |
| *col1a1b* | *dmh29* | p.(Gly1123Asp) | α1(I) p.(Gly958Asp) |
| *col1a2* | *dmh15* | p.(Gly882Asp) | α2(I) p.(Gly802Asp) |
| Knockout alleles in genes involved in Type I collagen processing |  |  |  |
| *bmp1a* | *sa2416* | p.(Arg522X) |  |
| *plod2* | *sa1768* | p.(Tyr679X) |  |

Mutant alleles are listed, along with the effect of the mutation on protein level (p-notation). For alleles causing an amino-acid substitution, the corresponding effect on the homologous human protein is listed in p-notation (35).

**Figure 1.**
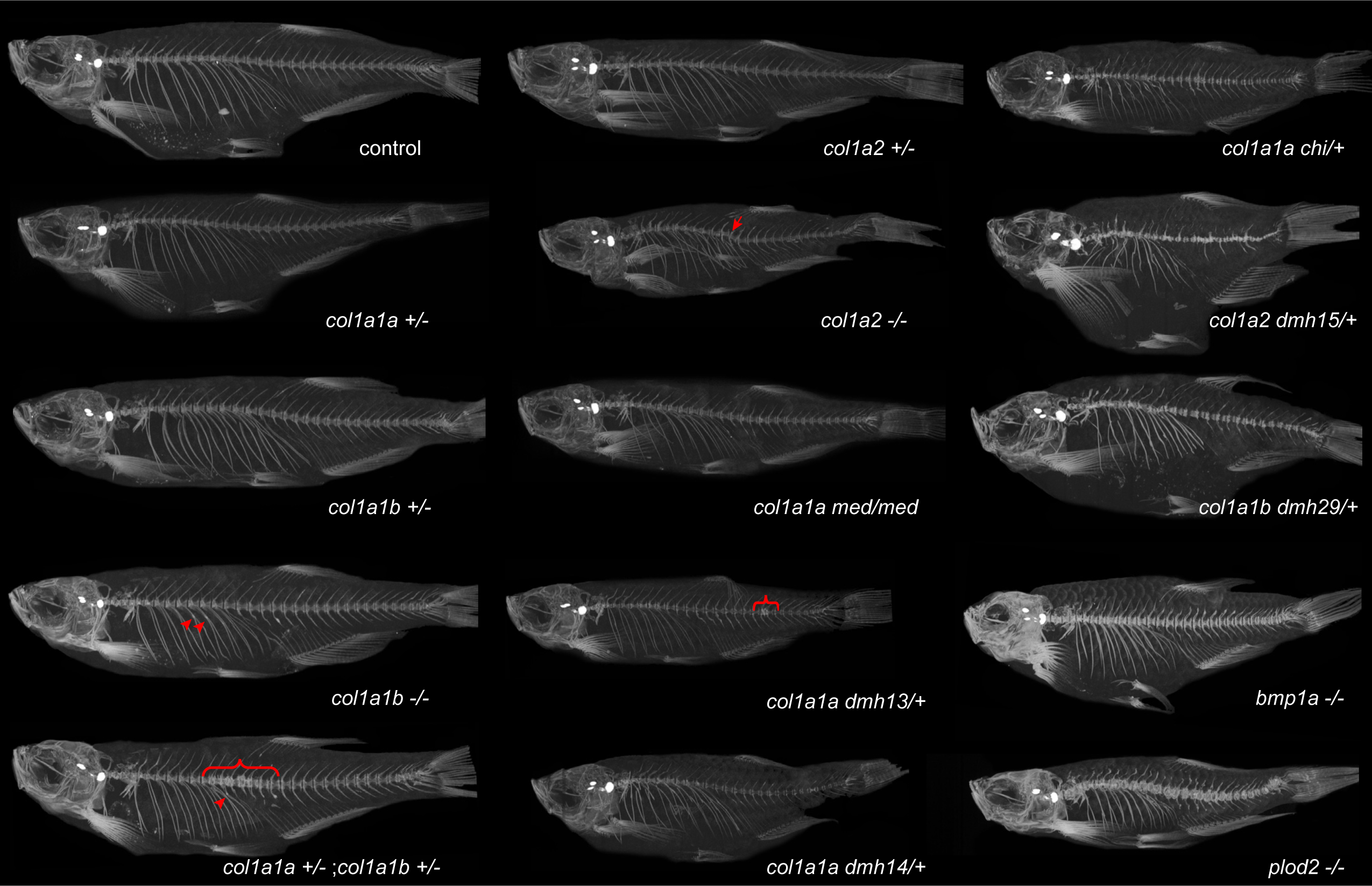
μCT images of different zebrafish models with affected type I collagen. For µCT scanning, we included several mutants with a non-sense or splice mutation in *col1a1a, col1a1b* or *col1a2*, generating a knockout of these genes (*col1a1a*^*−/−*^ not viable). Another set of mutants carries mutations resulting in substitutions of a Gly residue in α1(I) (*col1a1a*^*dmh13/+*^, *col1a1a*^*dmh14/+*^*and col1a1a*^*chi/+*^), α2(I) (*col1a2*^*dmh15/+*^) or α3(I) (*col1a1b*^*dmh29/+*^). *The microwaved* (*col1a1a*^*med/med*^) mutant carries a homozygous Glu substitution in α1(I). Finally, we included two mutant models with a knockout mutation in the *bmp1a* and *plod2* genes. Representative fish from each mutant genotype are shown. Callus formation in ribs (arrows), local compressions of the vertebral column (brackets), and kyphosis (arrowhead), are indicated (also listed in Supplementary Table 1).

**Figure 2.**
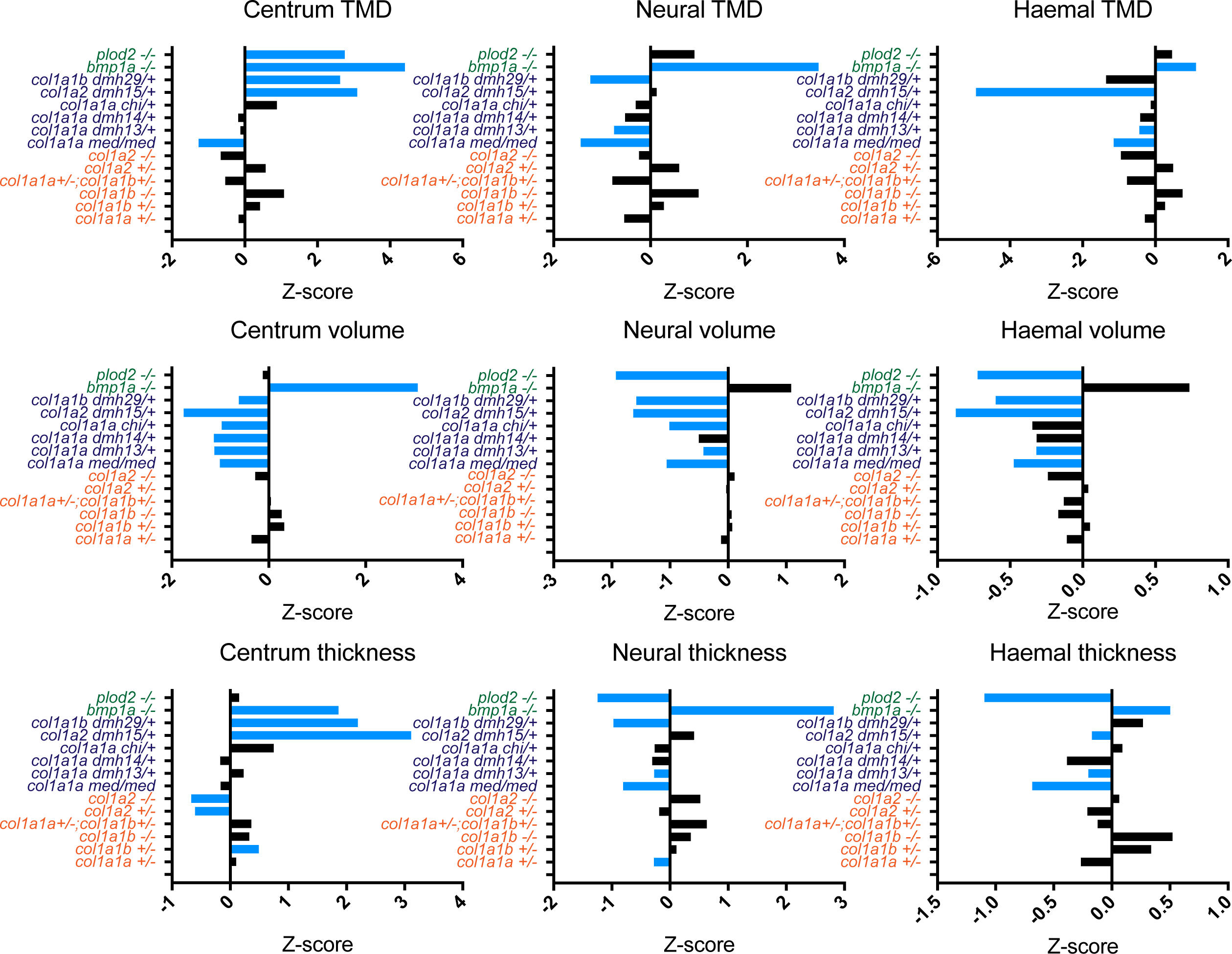
Inter- and intra-genotype phenotypic variability demonstrated by quantitative μCT analysis in FishCuT. **A.** For a set of parameters that quantify morphology and mineralization in the vertebral column of the different mutant populations, Z-scores are given for each mutant genotype (Z-score = ((mean value mutant – mean value control)/SD Control). On the Y-axis, mutants associated with a collagen processing defect are indicated in green, mutants with a qualitative defect in type I collagen are indicated in dark blue, and mutants with a quantitative defect in type I collagen are indicated in orange. Blue bars indicate values that were statistically significantly altered for that mutant genotype, compared to its control as analyzed by the global test (see supplementary figures 1-15). Note the large variability in the effect of each mutation on the different parameters.

### Heterozygous loss of both α1(I) and α3(I) in zebrafish is reminiscent of mild OI type I

Human patients with a heterozygous *COL1A1* null allele *(COL1A1* haplo-insufficiency) develop OI type I, characterized by mild bone fragility, relatively few fractures, and minimal limb deformities. In zebrafish two orthologues of the human *COL1A1* exist, namely *col1a1a* encoding α1(I) and *col1a1b* encoding α3(I). Two zebrafish knockout mutants with a premature stopcodon mutation in either *col1a1a* or *col1a1b* show absence of respectively α1(I) and α3(I) in the vertebral bone (Figure 3A). However, both qualitative and quantitative assessment of µCT scans of the vertebral column of heterozygous *col1a1a*^+/−^ or *col1a1b*^+/−^ zebrafish mutants revealed no skeletal abnormalities (Figure1, Supplementary Figure 1 and 2), which is most likely related to functional redundancy between both paralogs. Hence, we generated a double heterozygous knock-out mutant (*col1a1a*^+/−^;*col1a1b*^+/−^), which displays a mild skeletal phenotype, with evidence of spontaneous fractures (calluses in the ribs) and localized compression and mild malformation of the vertebral bodies in some of the mutant fish (Figure 1, Figure 4A). Quantitative measures of bone in this mutant revealed localized reduction of centrum volume, length and TMD (Supplementary Figure 3). However, these results were not found to be statistically significant, most likely due to the variable intra-genotype phenotypic penetrance, which is further demonstrated by the large spread of values for the different skeletal parameters in the group of mutant fish (Figure 4B).

**Figure 3.**
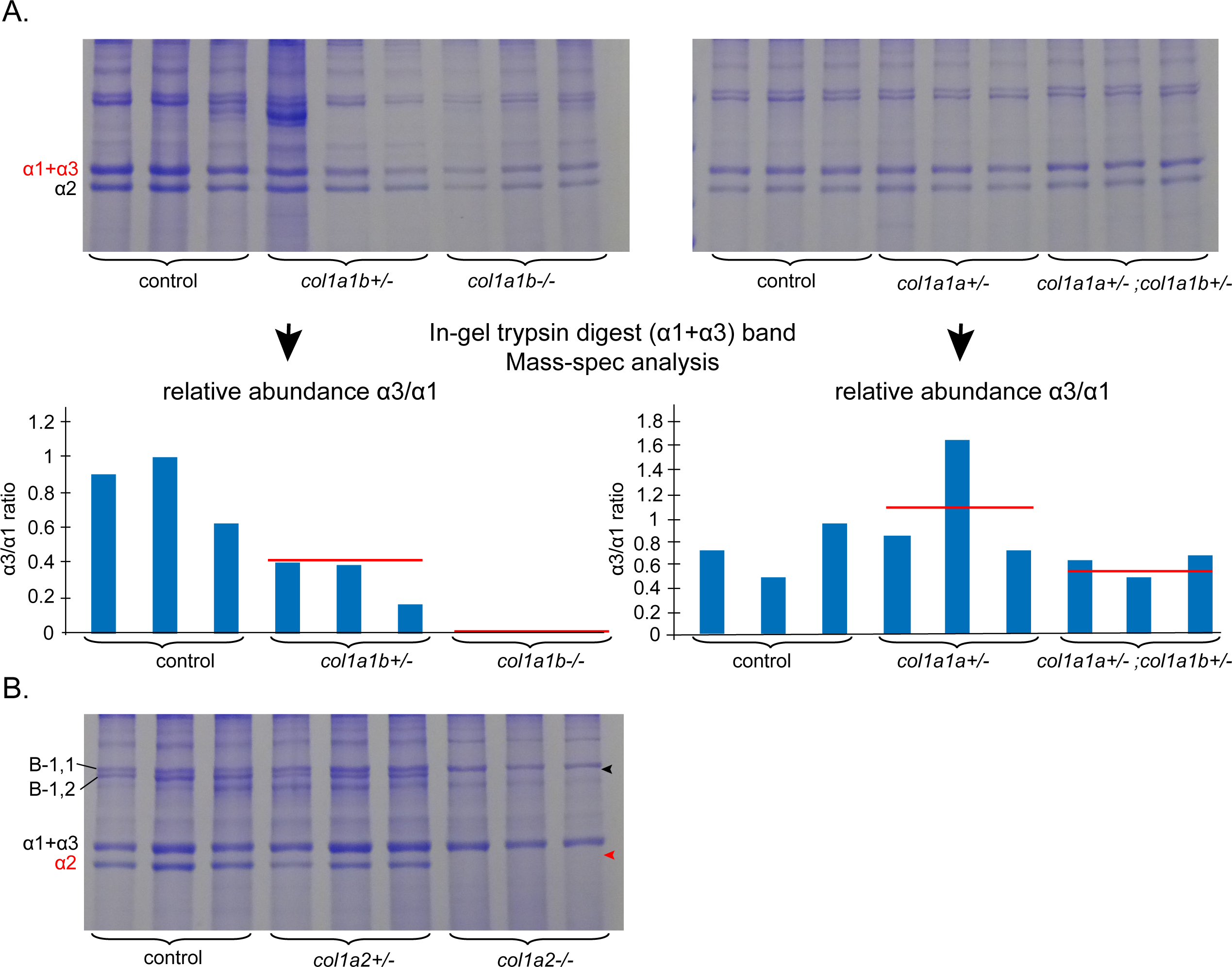
SDS-PAGE and mass-spectrometry analysis of bone collagen from adult spine of *col1a1a, col1a1b* and *col1a2* mutant fish with a quantitative type I collagen defect. **A.** SDS-PAGE and mass-spectrometry analysis of bone collagen from adult spine of *col1a1a, col1a1b* and *col1a2* mutant fish with a quantitative type I collagen defect. A. SDS-PAGE analysis of bone collagen extracted from individual adult spines of control fish, *col1a1b*^*+/−*^*, col1a1b*^*−/−*^*, col1a1a*^*+/−*^ and *col1a1a*^*+/−*^*;col1a1b*^*+/−*^ mutant fish. For each genotype three biological replicates were taken into account. Following SDS-PAGE separation, an in-gel trypsin digest of the α1(I)+α3(I) bands of each sample was performed and mass-spectrometry analysis was used to determine the relative amount of α1(I) and α3(I) in each sample. The expected α3(I)/α1(I) ratios (red lines in diagram) were approximated in samples from individual mutants (blue bars), illustrating that the reported non-sense mutations in *col1a1a* and *col1a1b* result in loss of the α1(I) chain and the α3(I) chain respectively. Accordingly, in *col1a1b*^*−/−*^ mutants, no tryptic peptides of α3(I) could be detected, confirming decay of mutant *col1a1b* mRNA transcripts. B. SDS-PAGE analysis of bone collagen extracted from individual adult spines of *col1a2*^*+/−*^ and *col1a2*^*−/−*^ mutant fish and control siblings. For each genotype three biological replicates were taken into account. *Col1a2*^*−/−*^ mutant fish show a complete absence of *col1a2* encoded α2(I) (red arrowhead). Note also the absence of the β1,2 dimers (black arrowhead) in *col1a2*^*−/−*^ mutant fish.

**Figure 4.**
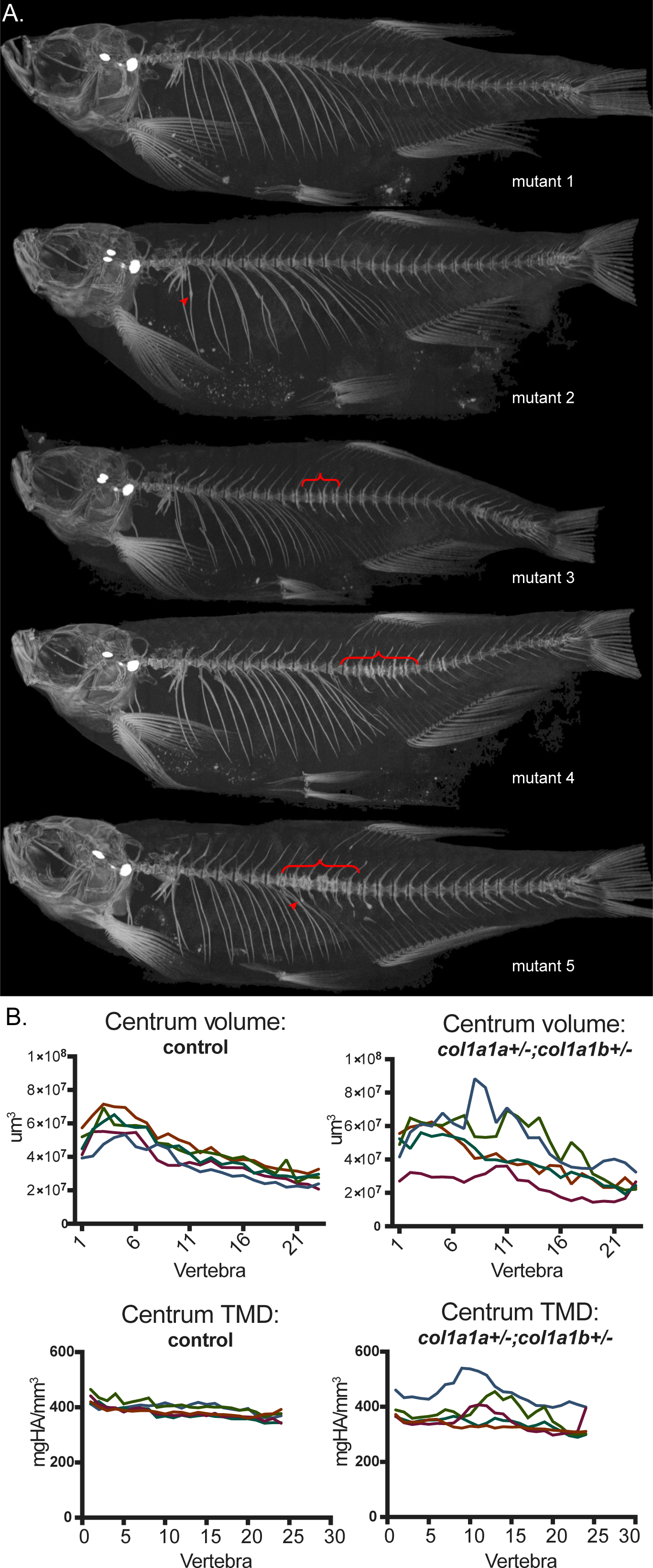
*col1a1a*^*+/−*^;*col1a1b*^*+/−*^ mutant fish exhibit intra-group variability in phenotypic penetrance. **A.** Different *col1a1a*^*+/−*^*;col1a1b*^*+/−*^ mutant siblings were analyzed using µCT scanning at 5 months of age (mutant 1- mutant 5). Upon comparison of the different mutant siblings, a variable penetrance of features indicative of compromised bone quality was observed, which include fractures (arrowheads) and local compressions in the vertebral column causing a distorted shape of the vertebral bodies (brackets) **B.** Intra-group variability for *col1a1a*^*+/−*^*;col1a1b*^*+/−*^ mutant fish is demonstrated for volume and TMD of the vertebral centra, with values in the mutant group showing a much larger spread than compared to the control group. The different colors in one graph represent different individual fish of the same group (mutant siblings or control siblings).

### A complete loss of zebrafish α1(I), but not α3(I), causes lethality in early larval stages

We next assessed the effect of a complete (homozygous) loss of α1(I) or α3(I) on survival and on skeletal integrity. In progeny resulting from an in-cross of *col1a1a*^*+/−*^;*col1a1b*^*+/−*^ mutant fish, all possible genotypes were shown to be present at 7 dpf, while from 15 dpf on some genotypes were underrepresented according to Mendelian predictions or even lost (Figure 5A). Eventually, all genotypes containing a homozygous knockout of *col1a1a* (loss of α1(I)) were found to be lethal by the age of 3 months. Homozygous mutant *col1a1b*^*−/−*^ mutants (loss of α3(I)) were present by the age of 3 months, however at reduced numbers if combined with heterozygous loss of *col1a1a* (*col1a1a*^*+/−*^;*col1a1b*^*−/−*^). Fish with a complete loss of α3(I), but intact α1(I) did not show significant alterations of bone mineralization or morphology (Supplementary Figure 4), although some fish showed evidence of fractures (Figure 1) (and increased TMD throughout the vertebral column, indicating an impaired bone quality.

**Figure 5.**
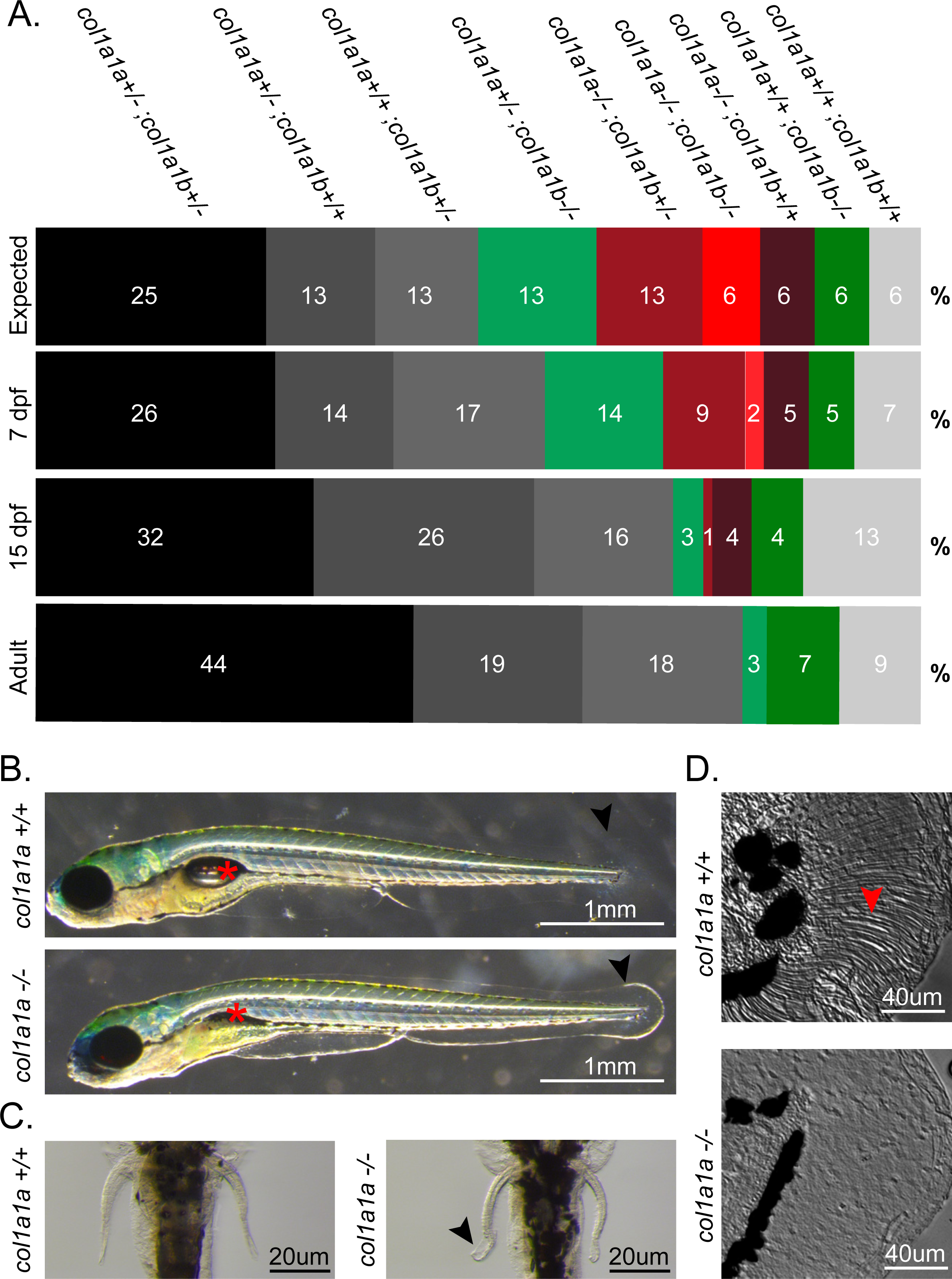
In-depth analysis of mutants with partial or complete loss of *col1a1a* and/or *col1a1b*. **A.** segregation analysis of the different mutant genotypes generated from an in-cross of *col1a1a*^*+/−*^*;col1a1b*^*+/−*^ double heterozygous mutant fish. Progeny was genotyped at 3 different time points (100 fish per time point): 7 dpf, 15 dpf and adult stage (3 months). The relative abundance of each genotype is given for each time point as a percentage. ‘Expected’ indicates the theoretical abundance of each genotype, following Mendelian segregation. All combinations with a complete loss of *col1a1a* (*col1a1a*^*−/−*^, indicated in red shades), are less viable and completely lost when progeny reaches adulthood. Genotypes which contain a complete loss of *col1a1b* (*col1a1b*^*−/−*^, indicated in green shades) are still present at adult stage. Black indicates the parental genotype, white indicates wild type fish, and grey represents the other remaining genotypes. **B.** Phenotype of *col1a1a*^*−/−*^ mutant larvae at 7 dpf, compared to wild type siblings. *Col1a1a*^*−/−*^ mutant larvae present with a non-inflated swimbladder (asterisk) and a ruffled finfold (arrowhead). **C.** Ventral view and detail of the pectoral fins showing frilly distal margins (arrowhead) in the pectoral fins of *col1a1a*^*−/−*^ mutant larvae at 7 dpf. **D.** Detail of the *col1a1a*^*−/−*^ mutant larvae finfold at 7 dpf, showing an absence of fin actinotrichia fibers, which can be clearly observed in the finfold of wild type siblings (arrow).

Since the knockout of *col1a1a* compromises viability from 7dpf on, the general morphology of *col1a1a*^−/−^ mutant larvae was assessed at this stage. These larvae lacked the presence of an inflated swim bladder, a structure which is essential for survival once larvae are required to actively feed (Figure 5B). Furthermore, the distal margins of the pectoral fins, and the finfold, were shown to be ruffled in these mutants (Figure 5B-C). These fin structures are structurally supported by fin actinotrichia, which are pre-fin-ray structures, composed of collagen fibers (31). Upon detailed examination of the finfold, these actinotrichia were found to be absent in *col1a1a*^*−/−*^ mutant larvae (Figure 5D). This is in concordance with earlier studies reporting these structures to only contain type I collagen as α1(I) homotrimers (31, 32), while in other tissues both α1(I) and α3(I) are present (31, 33).

### Complete loss of α2(I) in zebrafish leads to soft connective tissues abnormalities, reminiscent of the human Ehlers-Danlos syndrome

We studied a zebrafish mutant with a splice mutation in *col1a2* resulting in absence of α2(I) (Figure 3B). In human, heterozygosity for *COL1A2* null mutations (COL1A2 haplo-insufficiency) is not known to be pathogenic, while a complete absence of α2(I) chains is associated with the cardiac-valvular type of EDS, a soft connective tissue disorder characterized by skin fragility, joint hypermobility and early-onset, severe and progressive cardiac valvular defects without bone involvement (6). Similarly, zebrafish with a heterozygous *col1a2* knockout displayed no morphological or skeletal abnormalities (Figure 6A, Supplementary Figure 5), while homozygous *col1a2* knock-outs were severely affected with marked kyphosis of the vertebral column at the level of transitioning pre-caudal to caudal vertebrae (Figure 1, Figure 6A arrowhead). However, no other obvious skeletal abnormalities could be detected in these mutants, and only minor and non-significant effects on mineralization and morphology in the vertebral column of *col1a2*^*−/−*^ mutant fish were observed (Supplementary Figure 6).

**Figure 6.**
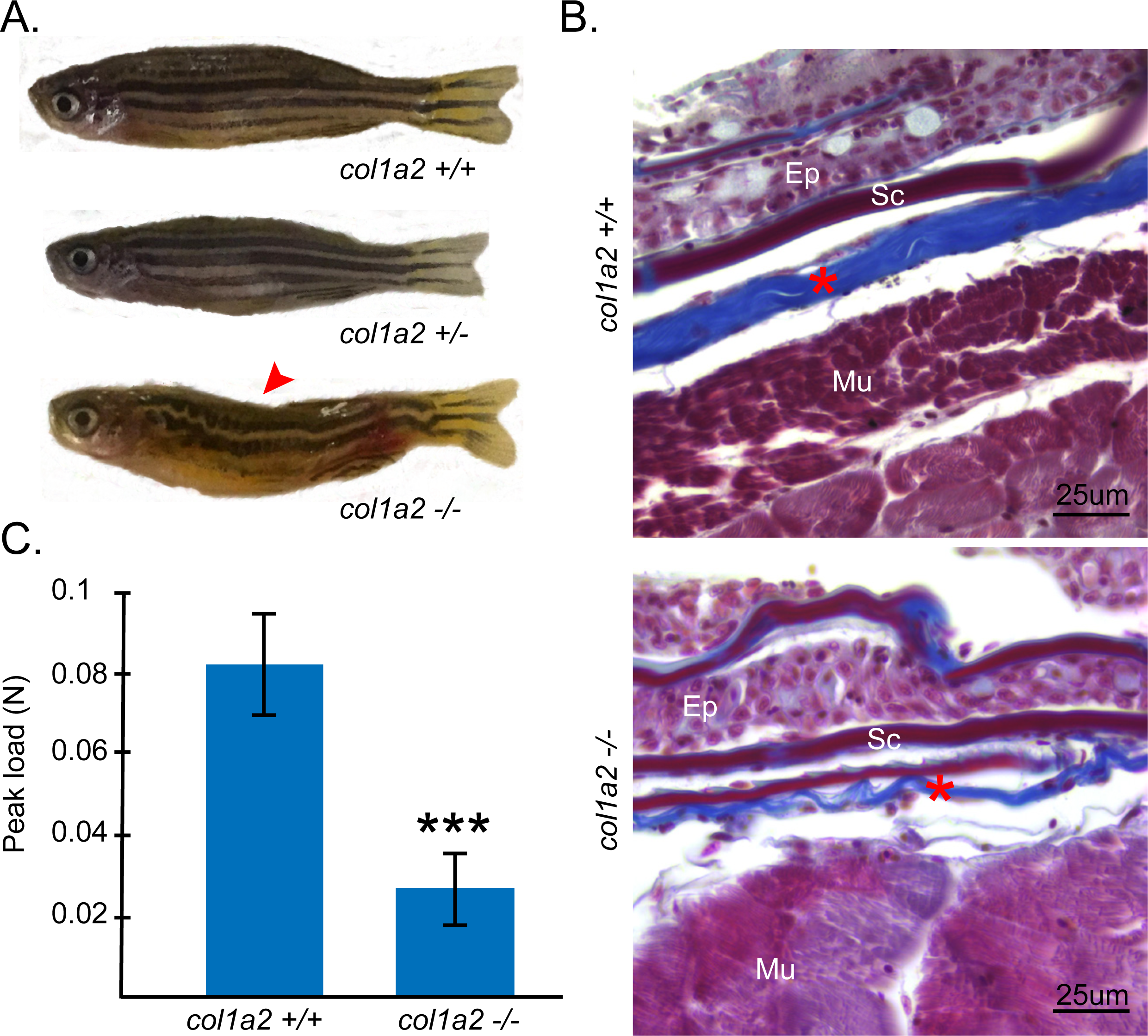
In-depth analysis of *col1a2*^*+/−*^ and *col1a2*^*−/−*^ mutant fish. **A.** Visual phenotype of *col1a2* mutant and wild type siblings. Both *col1a2*^*+/+*^ control, and *col1a2*^*+/−*^ mutant fish display a normal morphology, while *col1a2*^*−/−*^ mutant are dysmorphic and display kyphosis (arrowhead). The typical stripe pattern in the skin is disturbed in *col1a2*^*−/−*^ mutant fish, but normal in *col1a2*^*+/+*^ controls and *col1a2*^*+/−*^ mutant fish. **B.** Masson’s Trichrome staining of histological sections of the skin of an adult *col1a2*^*−/−*^ mutant fish and a *col1a2*^*+/+*^ control sibling. Note the much thinner dermis (asterisk), composed of layers of collagen fibrils (stained blue) in *col1a2*^*−/−*^ mutant fish when compared to *col1a2*^*+/+*^ controls. **C.** Average maximum tensile strength, measured by biomechanical testing of skin flaps dissected from 5 adult *col1a2*^*−/−*^ mutant fish and 5 *col1a2*^*+/+*^ control fish, illustrating strongly decreased strength of soft connective tissues in mutant fish (p<0.0001). Other abbreviations: Ep, Epidermis; Mu, Muscle fibers; Sc, Scale.

Given the soft connective tissues involvement in human patients with a complete loss of α2(I) (6), we explored the presence of similar defects in *col1a2*^*−/−*^ mutant fish. The skin of *col1a2*^*−/−*^ mutant fish was more fragile and easily damaged, compared to the skin of wild type control fish. Histological analysis of adult skin showed that the dermis, which is composed of two collagen fibril layers in zebrafish, is half the thickness in mutant fish when compared to dermis of wild type siblings (Figure 6B). Biomechanical load-to-failure strength of soft connective tissues was analyzed by determining the ultimate tensile strength of tissue specimens from 5 *col1a2*^*−/−*^ mutant and 5 control fish (Figure 6C). Tissues lacking α2(I) ruptured at a significantly lower peak load (0.027 ± 0.0079 N), when compared to tissue samples from wild type siblings (0.081 ± 0.011 N, p<0.0001), indicating significantly diminished strength of soft connective tissue in *col1a2*^*−/−*^ mutant fish. Histological sections of the adult zebrafish heart from *col1a2*^*−/−*^ mutants showed normal morphological appearances of the cardiac valves and *col1a2*^*−/−*^ larvae displayed normal cardiac function and blood flow (results not shown).

### Genotype/Phenotype relations in mutants with impaired quality of type I collagen

We next studied the skeleton of a set of mutants carrying a point mutation in either of the type I collagen encoding genes, leading to impaired type I collagen quality (Figure 1, Table 1, Supplementary Figures 7-13).

The *col1a2*^*dmh15/+*^ mutant represents the most detrimental phenotype of all analyzed mutants (Figure 1, Supplementary Figure 12), with heavily distorted, misshapen and overmineralized axial and cranial skeletons. This is in concordance with human data as Gly substitutions in the homologous region of α2(I) are associated with lethal OI in human patients (Figure 7)(34). The *col1a1a*^*chi/+*^ mutant, reported earlier (21, 35), carries the same mutation that was identified in a human patient with OI type III (34), and displays a moderate to severe phenotype with skeletal malformation including shorter vertebral bodies and evidence of fractures (Figure 1, Supplementary Table 1, Supplementary Figure 9). The alleles *dmh13, dmh14* and *dmh29*, all cluster in the major ligand binding region 3 (MLBR3) of type I collagen (Figure 7), and while this region represents a hotspot for ligand interactions, mutations in this region are associated with variable phenotypic outcomes (3, 34). This was also the case when comparing the three different mutants, with *col1a1b*^*dmh29/+*^ fish displaying the most severe skeletal phenotype with kyphoscoliosis and shorter, thicker and overmineralized vertebral bodies and frequent rib fractures (Figure 1, Supplementary Table 1, Supplementary Figure 10, 11 and 13). The *col1a1a med* allele, reported earlier by Asharani et al. (20), was shown to have no effect in heterozygous state but caused pronounced effects on TMD, volume and thickness of the vertebral bodies when present in homozygous state (Supplementary Figure 7-8). However, as no comparable mutations have been identified in human patients this *col1a1a*^*med/med*^ mutant is likely less relevant for human OI.

**Figure 7.**
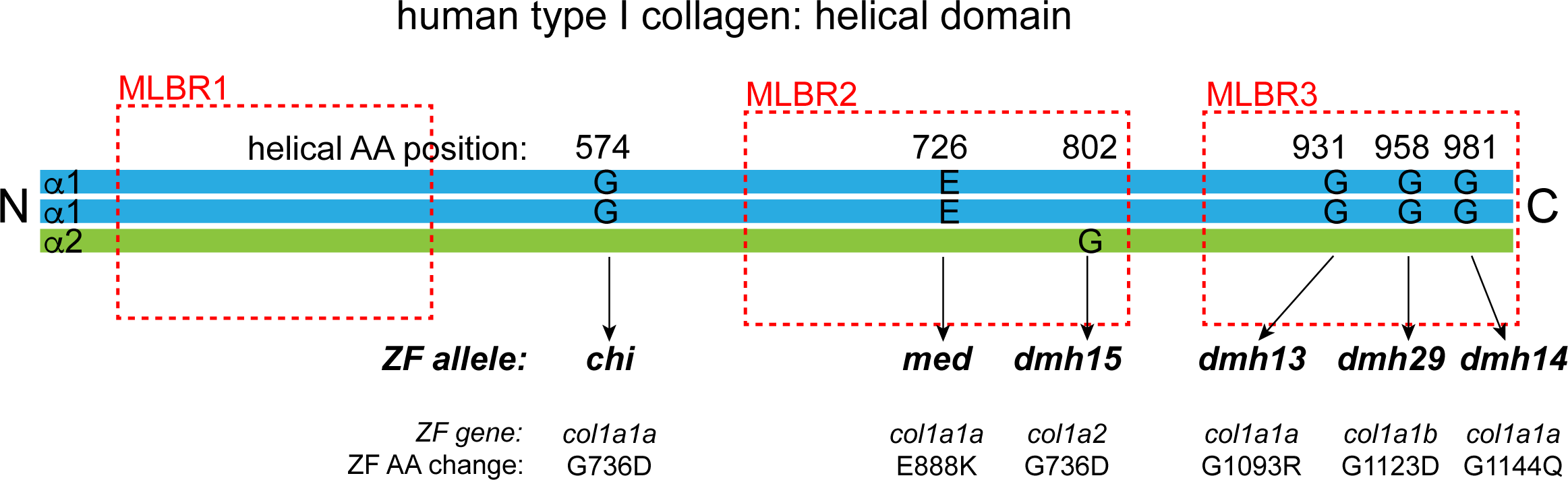
Zebrafish alleles representing amino-acid substitutions in type I collagen mapped on the helical domain of human type I collagen. The α1 and α2 chains of the human type I collagen are schematically depicted with the three major ligand binding regions (MLBR), indicated with red stripped lines (adapted from Sweeney et al, (35). The homologous positions to the zebrafish (ZF) alleles (as determined by Clustal W analysis), representing mutations that cause an amino-acid (AA) substitution, are indicated with arrows. The numbering of the zebrafish alleles indicates the total protein postion of the residue.

To detect the presence of collagen overmodification, which is typically caused by collagen I glycine substitutions in human OI patients, collagen from adult bone was extracted by heat denaturation and subjected to SDS-PAGE. While migration of the α-chains was not affected in bone from *col1a2*^*dmh15/+*^, *col1a1b*^*dmh29/+*^ and *col1a1a*^*med/med*^ mutants, retarded α-chain mobility, indicating collagen overmodification, was observed for *col1a1a*^*chi/+*^, *col1a1a*^*dmh13/+*^ and *col1a1a*^*dmh14/+*^ mutants (Figure 8).

**Figure 8.**
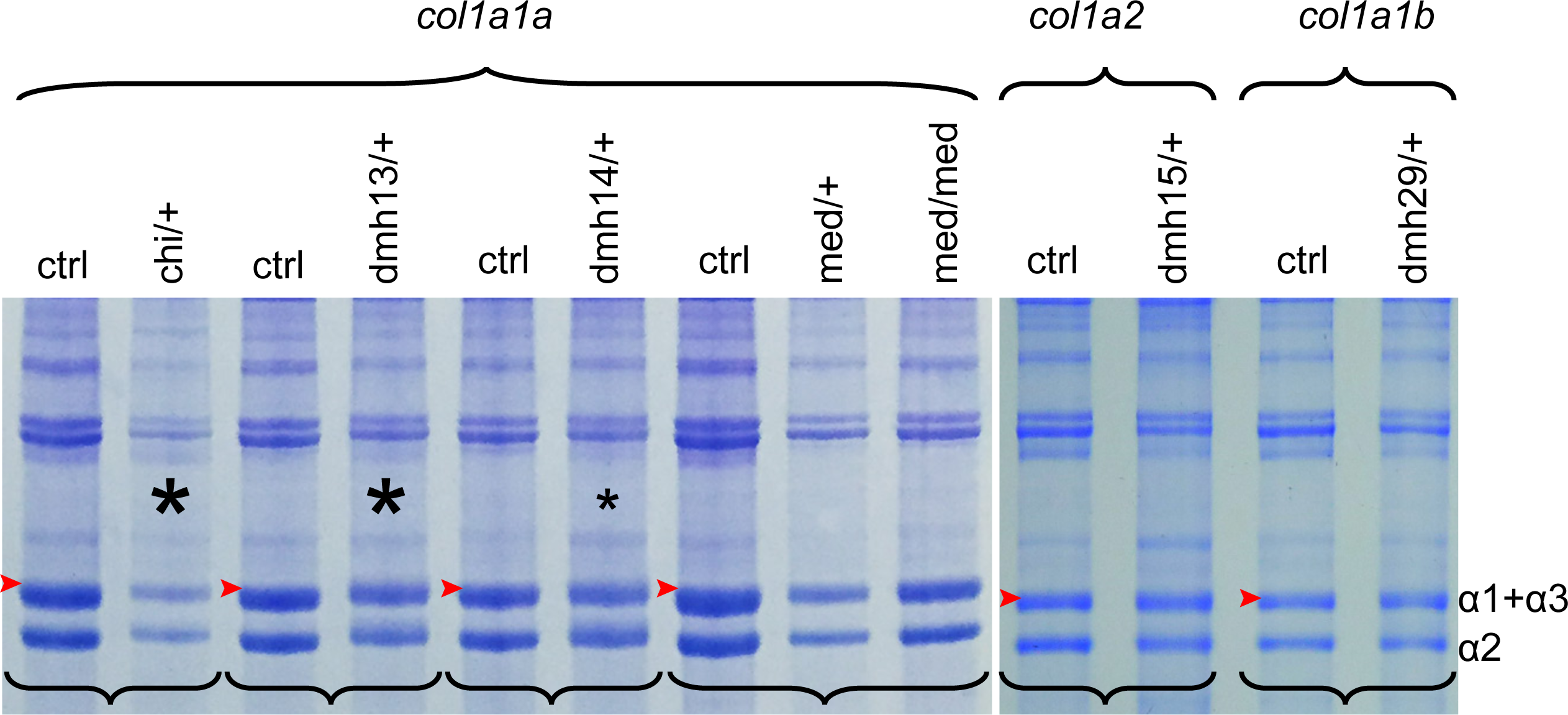
SDS-PAGE of collagen extracted from the bone of adult mutants with qualitative type I collagen defects. SDS-PAGE analysis of acid soluble collagen extracts from different mutant genotypes carrying amino-acid substitutions in α1(I), α2(I) or α3(I), and from matching wild type control (ctrl) siblings (sibling pairs indicated by brackets) indicate that alpha band migration is markedly slower for *col1a1a*^*chi/+*^ and *col1a1a*^*dmh13/+*^ mutants, and only slightly decreased in *col1a1a*^*dmh14/+*^ mutants (lanes indicated by asterisks), when compared to the corresponding control. No apparent signs of collagen overmodification could be detected for the other mutant genotypes.

## DISCUSSION

In this work, we present a skeletal phenomic analysis of a set of zebrafish with mutations in type I collagen that model different type I collagenopathies, including the classical types of OI and a specific subtype of EDS. By systematically analyzing the skeletal phenotypes in this set of mutants we illustrate the high ability of zebrafish mutants to reproduce the genetic and phenotypic features of type I collagenopathies.

OI in human patients is characterized by bone fragility, leading to a higher susceptibility to fractures, and in addition, misshapen bones and skeletal malformation can be observed (7). The presence of these disease features and the extent to which they manifest is largely associated with the specific subtype of OI (7). As such, quantitative defects in type I collagen, underlying OI type I, tend to cause mild skeletal defects in human patients with an increased susceptibility to fractures but minimal bone deformity. Zebrafish mutants with similar quantitative type I collagen defects (*col1a1a*^*+/−*^;*col1a1b*^*+/−*^ mutants) also presented with increased fracture risk, apparent from the presence of spontaneous fractures in the ribs, while skeletal deformities were minimal with only localized and mild compressions of the vertebral column in some of the mutant fish. Conversely, qualitative type I collagen defects in zebrafish evoked a more pronounced effect on the skeleton, with frequent fractures, kyphoscoliosis and malformation of the vertebral bodies, and misshapen ribs and fin bones. These phenotypes correspond to the moderately severe, progressive deforming or lethal clinical phenotypes of OI type II-IV. Importantly, the more severely affected mutants of this group (*dmh15* and *dmh29*) also displayed overmineralization in the vertebral column, a feature often seen in human OI, and especially associated with the more severe cases in human patients (36, 37). Additionally, biochemical analysis argued for type I collagen overmodification, one of the typical features in human OI, in some of the mutants with a qualitative defect. Moreover, the variability of the excessive posttranslational modification throughout this set of mutants corresponds to the situation in human patients with OI type II-IV, where variable collagen overmodification has been related to the position of the mutation in the type I collagen helix (34, 38).

A hallmark feature of OI in humans is the large clinical spectrum of phenotypes, with disease presentation that can differ from very mild to perinatal lethal, even with the same causative mutation. The underlying molecular basis of this variability remains largely not understood. The zebrafish mutants described here similarly show significant variation in their expressivity. Intra-genotype variability is illustrated in *col1a1a*^*+/−*^;*col1a1b*^*+/−*^ mutant siblings, as some show fractured ribs and/or vertebral compressions, while others do not. In addition, there is a large variability in vertebral morphology and mineralization in mutant siblings, in contrast to control siblings. A likely explanation for the intra-genotype phenotypic variability in zebrafish mutants is the highly polymorphic nature of its genome, which closely resembles human genetic variability. This makes them particularly useful to unveil genetic determinants that can be used to map potential regulation of phenotypic variability in humans. This is less obvious in mouse models where mostly inbred strains are used for disease modeling (39, 40).

In zebrafish, type I collagen has been shown to have a different composition than in mammals, with two orthologues of the human *COL1A1* gene, namely *col1a1a* and *col1a1b*, encoding the α1- and α3-chain respectively. Some insights in these differences have already been addressed previously (19), and by studying different knock-out mutants for the type I collagen genes we further extend this knowledge. Heterozygous loss of either α1(I) or α3(I) did not induce any skeletal abnormalities, while reduced levels of both α1(I) and α3(I) (*col1a1a*^*+/−*^; *col1a1b*^*+/−*^ mutants) causes a mild skeletal phenotype with increased bone fragility, arguing for interchangeability and functional redundancy between α1(I) and the homologous α3(I). Complete loss of the α1(I) chain causes lethality after 7 dpf while loss of α3(I) only shows increased early lethality if the amount of α1(I) is also diminished. This suggests a gene/protein dosage effect, where α1(I) is more abundant than α3(I), as hypothesized in our previous work (19). The functional similarity between α1(I) and α3(I) is further illustrated by the fact that glycine substitutions in both genes can cause severe skeletal phenotypes, arguing for a dominant negative effect on type I bone collagen and thus incorporation of a substantial amount of both α1(I) and α3(I) into the type I collagen triple helix. However, to obtain a full understanding of zebrafish type I collagen, the exact trimer stoichiometry in different tissues, and the abundance of each possible trimer composition should be examined, which is technically challenging due to the small specimen size of zebrafish compared to the large amount of input material needed for standard biochemical analysis techniques.

In humans, haploinsufficiency for the α2(I)-chain does not cause a clinically significant phenotype, while complete loss of α2(I) leads to the very rare cardiac valvular sub-type of the EDS, associated with soft connective tissue defects, including skin and joint manifestations and propensity to cardiac valvular defects, rather than apparent skeletal manifestations (6). These trends were recapitulated in zebrafish, as *col1a2*^*+/−*^ mutants are asymptomatic, whereas *col1a2*^*−/−*^ mutants display an EDS-like phenotype with defects in soft connective tissue. While µCT-scanning showed marked kyphosis in the vertebral column of *col1a2*^*−/−*^ mutants, the vertebral body morphology displayed no deviation from the normal hourglass shape. Therefore, the kyphosis seen in these mutants is likely related to defects in the fibrous joints between the vertebral bodies, concordant with the location of kyphosis being the area bearing the highest mechanical force in the fish vertebral column (34). Similar to these observations, patients with the same molecular defect display joint dislocations but no apparent bone phenotype (6). Interestingly, *col1a2*^*−/−*^ mutant fish showed fragile skin upon handling and histological analysis demonstrated a much thinner dermis in the skin of mutant fish when compared to wild type siblings. Consistent with these findings, bio-mechanical testing argued for a strongly reduced strength of soft connective tissues in *col1a2*^*−/−*^ mutant fish. This relates to other clinical hallmark features of the skin in human cardiac-valvular EDS patients (6). Preliminary experiments in *col1a2*^*−/−*^ mutant fish could not provide any proof of defects in cardiac morphology or functioning. However, more extensive analyses are needed to further explore a possible impaired cardiac function in a follow-up study, focusing on an in-depth characterization of this mutant, which was beyond the scope of the study presented here.

Taken together, we have analyzed a large set of zebrafish models with mutations in type I collagen that represent different type I collagenopathies, including the classical types of OI and a specific subtype of EDS, and illustrate the high phenotypic and genetic similarity of these zebrafish mutants with human type I collagenopathies. We further report the first zebrafish model of the human Elhers-Danlos syndrome, which harbors characteristic defects in the soft connective tissues. Our study surpasses the analysis at single mutant model level, but illustrates the occurrence of relevant phenotypic patterns and characteristics across a large set of genetically distinct zebrafish models. These results, taken together with the feasibility of zebrafish for easily accommodating large genetic and phenotypic screens, argue for zebrafish as a promising tool to further dissect the underlying basis of phenotypical variability in human type I collagenopathies, such as OI. Furthermore, we provide more insight into the biology of zebrafish type I collagen and its consequences, which are relevant in the study of any human type I collagen related disease by using zebrafish as a model.

## ACKNOWLEDGMENTS

This work was supported by the Ghent University Methusalem grant BOF08/01M01108 to ADP and by funding from the Belgian Science Policy Office Interuniversity Attraction Poles (BELSPO-IAP) program through the project IAP P7/43-BeMGI. RYK acknowledges NIH/NIAMS awards AR066061 and AR072199, DRE NIH/NIAMS awards AR037318 and AR036794, and MPH NIH award U01DE024434. The Zebrafish International Resource Center is supported by grant #RR12546 from the NIH-NCRR. FM is a Senior Clinical Investigator at the Fund for Scientific Research Flanders-Belgium. CG is supported by a postdoctoral fellowship from the Belgian American Educational Fund (BAEF) at the UW.

## Author contributions

Project design: AW, PJC, SS, ADP, FM, MPH, KH, SF and DRE. Data collection: CG, RYK, JWB, PV, HDS, MW, PS and BG. Data analysis: CG, RYK, MW. Data interpretation: CG, RYK, DRE, FM and AW. Drafting manuscript: CG and AW. Revising the manuscript content and approving final version of manuscript: all authors.

## Supplementary Table and Figure legends

**Supplementary Table 1.**
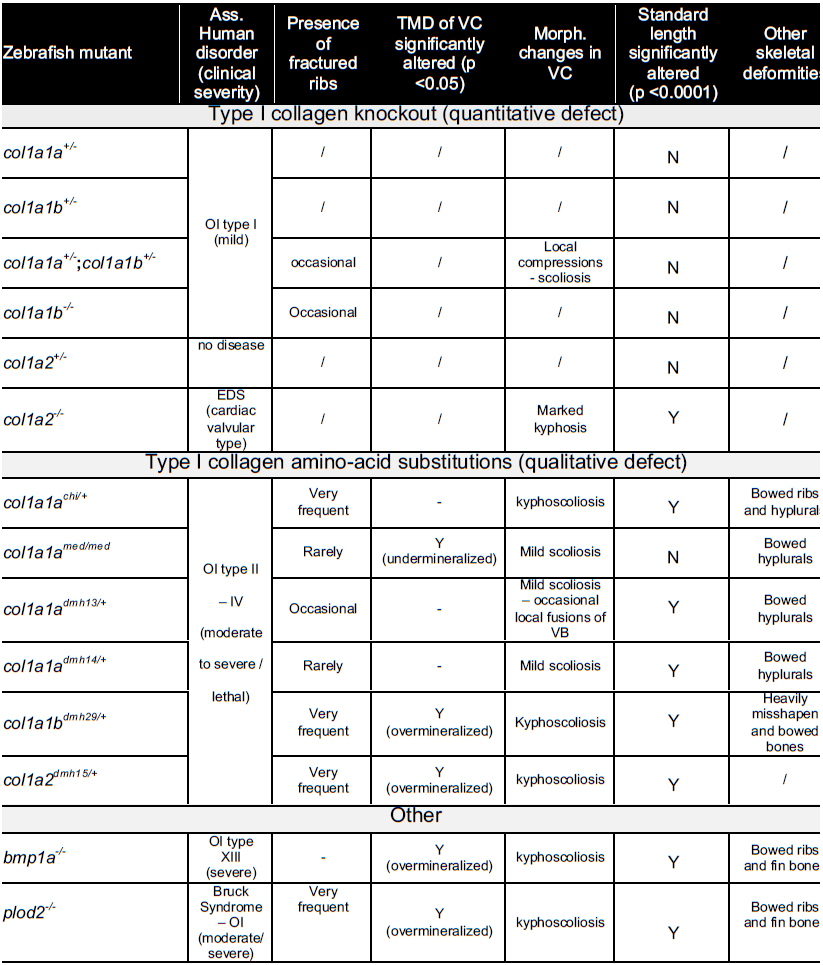
Skeletal features observed upon μCT-scanning of adult mutant fish included in this study. For each mutant model, the specific disorder and clinical severity associated with a similar mutation in human patients is listed. In addition, the presence of fractures, significant alterations of tissue mineral density (TMD) of the vertebral column (VC), morphological changes in the VC, and the presence of other skeletal deformities were assessed on reconstructed µCT scans and are listed for each mutant. Finally, significant alterations (p<0.001) of the mean standard length (SL) are indicated with yes (Y) or (N), see Supplementary Figure 16 for boxplot representation of SL measurements per genotype.

**Supplementary Figure 1.**
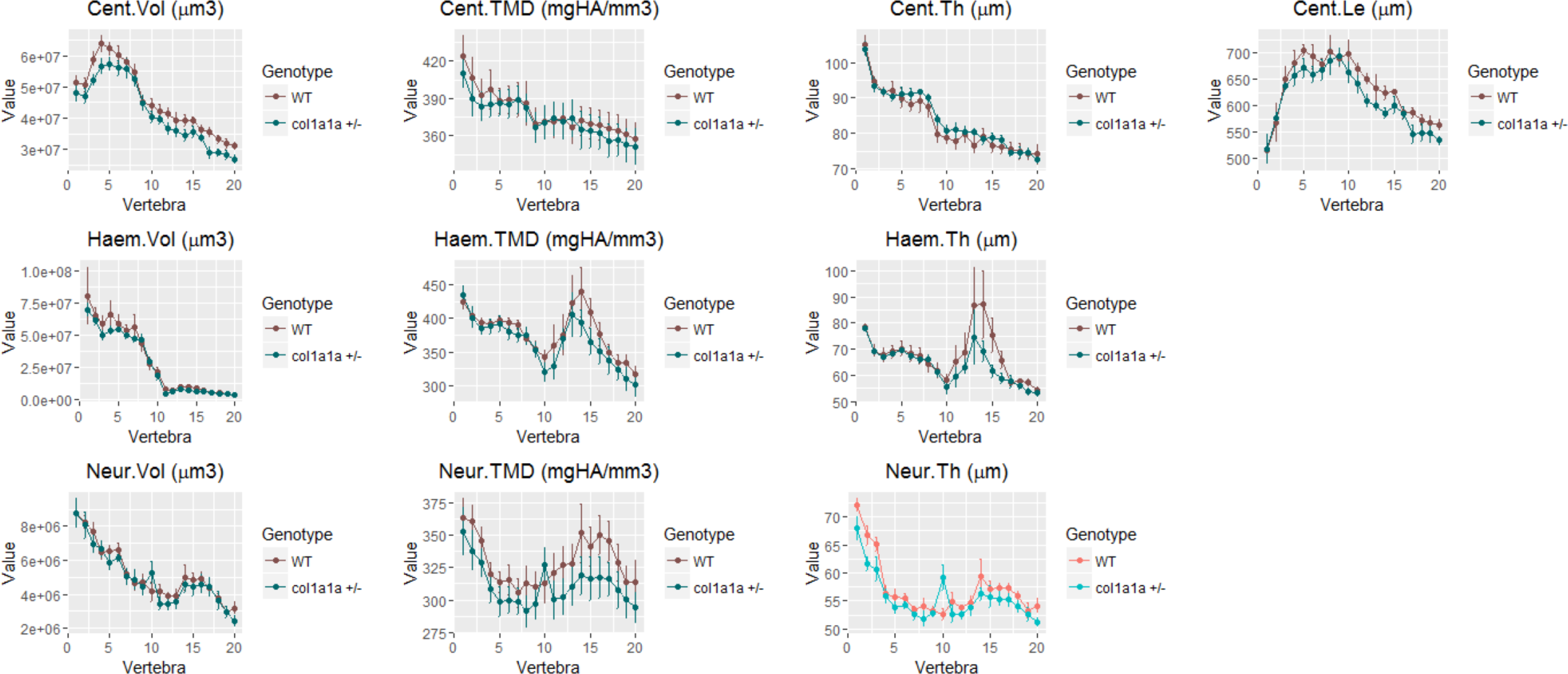
Quantitative µCT-scanning analysis of the vertebral column of *col1a1a*^*+/−*^ mutant fish, using FishCuT software. Whole-body µCT scans were acquired in *col1a1a*^*+/−*^ mutant adult fish and wild type control siblings. Individual vertebrae were isolated using FishCuT which segments each vertebral body into three skeletal elements: the Neural Arch (Neur.), Centrum (Cent.), and Haemal Arch/Pleural Ribs (Heam.). For each skeletal element of each vertebra, FishCuT computed the following parameters: Tissue Mineral Density (TMD), Volume (Vol), Thickness (Th.) and Length (Le., centrum only). For analysis, each combination of element/outcome is computed as a function of vertebra number, and subjected to the global test. Plots, generated in R statistics, associated with a significant difference in the mutant population, compared to wild type control siblings, are colored in a lighter coloring scheme

**Supplementary Figure 2.**
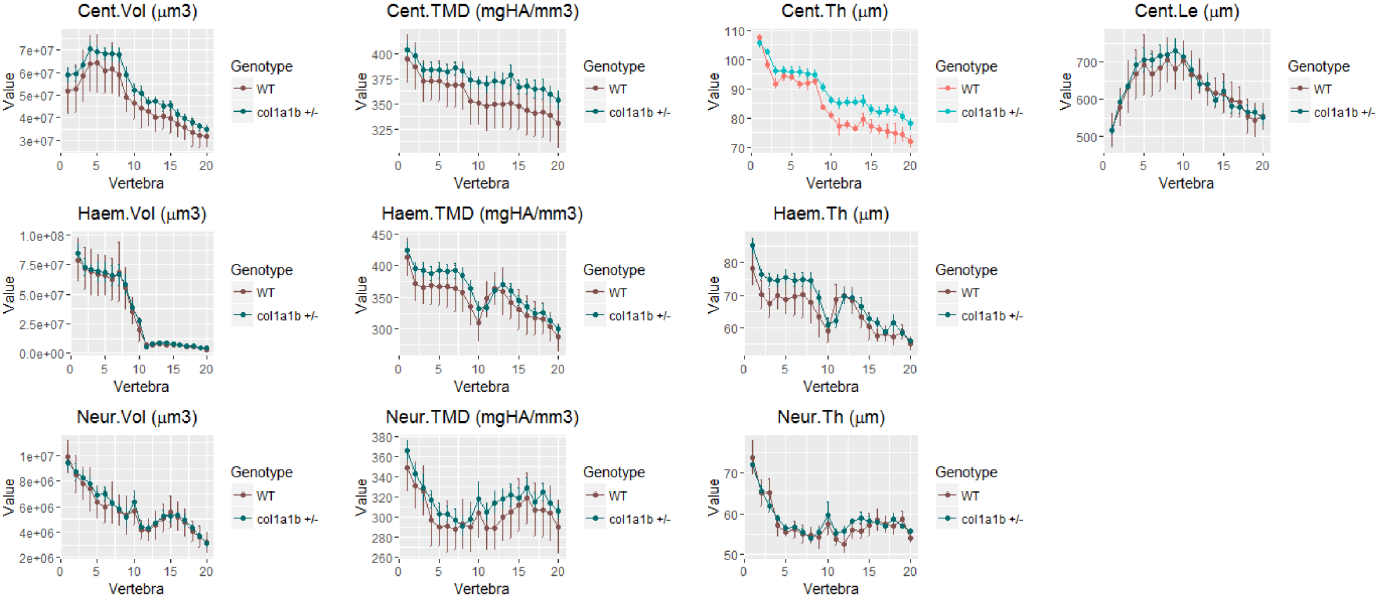
Quantitative µCT-scanning analysis of the vertebral column of *col1a1b*^*+/−*^ mutant fish, using FishCuT software. Whole-body µCT scans were acquired in *col1a1b*^*+/−*^ mutant adult fish and wild type control siblings. FishCuT analysis was performed and interpreted as explained previously (Supplementary Figure 1 legend).

**Supplementary Figure 3.**
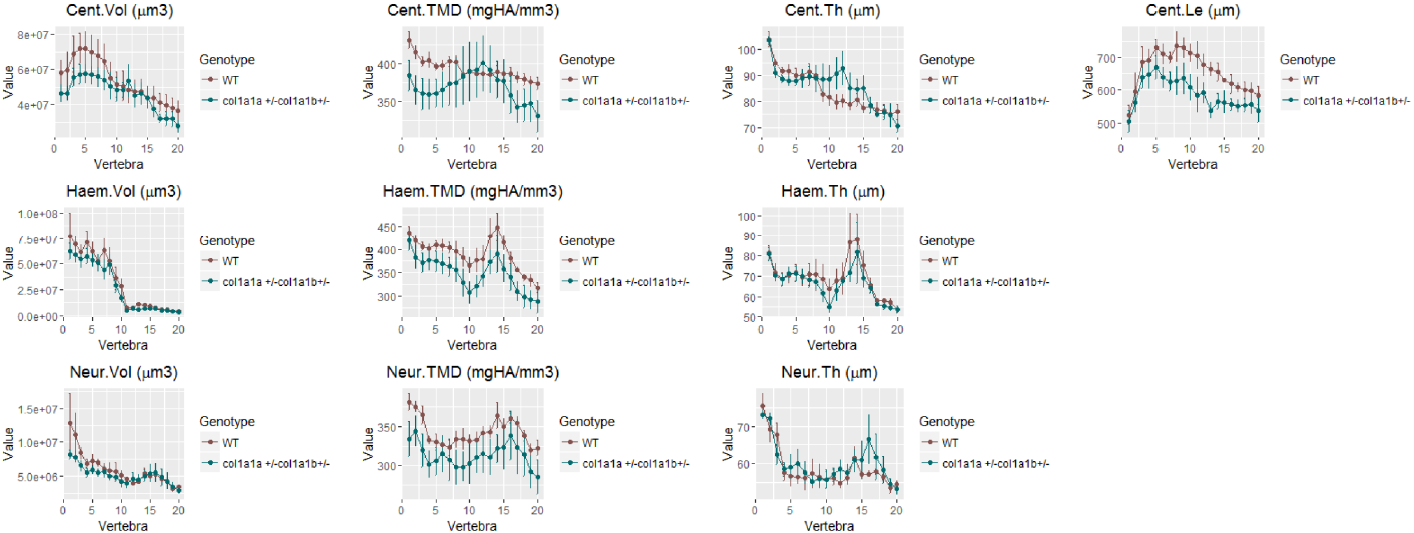
Quantitative µCT-scanning analysis of the vertebral column of *col1a1a*^*+/−*^*col1a1b*^*+/−*^ mutant fish, using FishCuT software. Whole-body µCT scans were acquired in *col1a1a*^*+/−*^*;col1a1b*^*+/−*^ mutant adult fish and wild type control siblings. FishCuT analysis was performed and interpreted as explained previously (Supplementary Figure 1 legend).

**Supplementary Figure 4.**
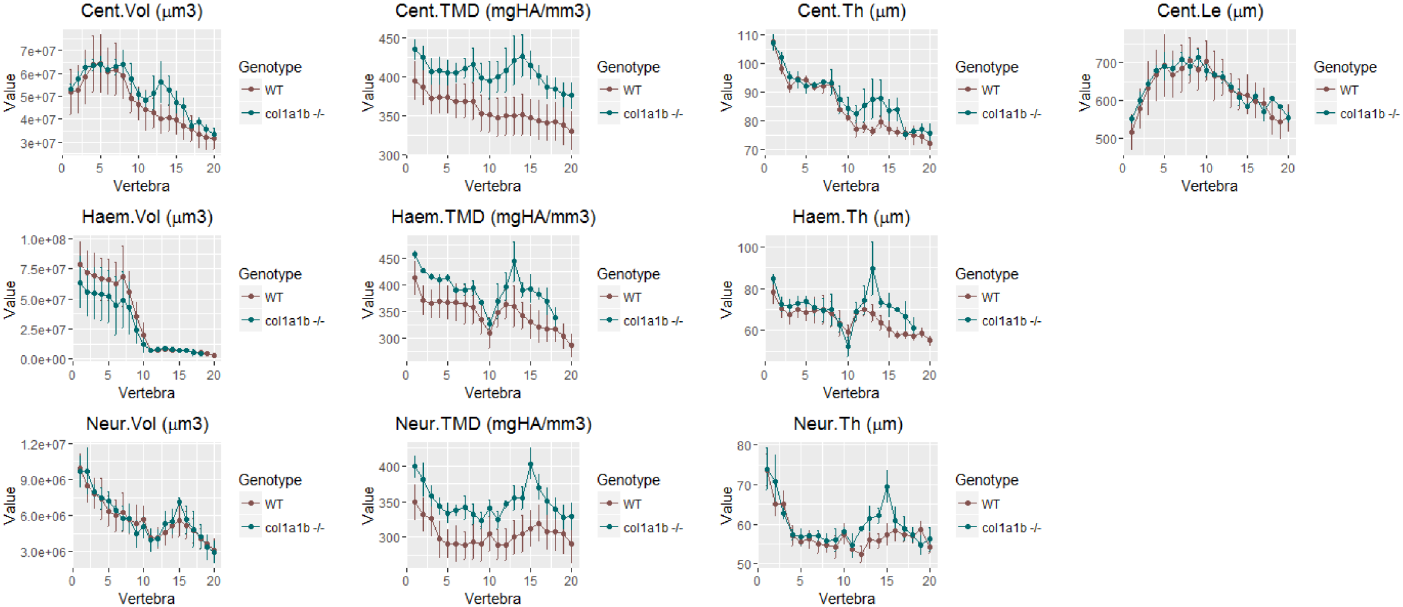
Quantitative µCT-scanning analysis of the vertebral column of *col1a1b*^*−/−*^ mutant fish, using FishCuT software. Whole-body µCT scans were acquired in *col1a1b*^*−/−*^ mutant adult fish and wild type control siblings. FishCuT analysis was performed and interpreted as explained previously (Supplementary Figure 1 legend).

**Supplementary Figure 5.**
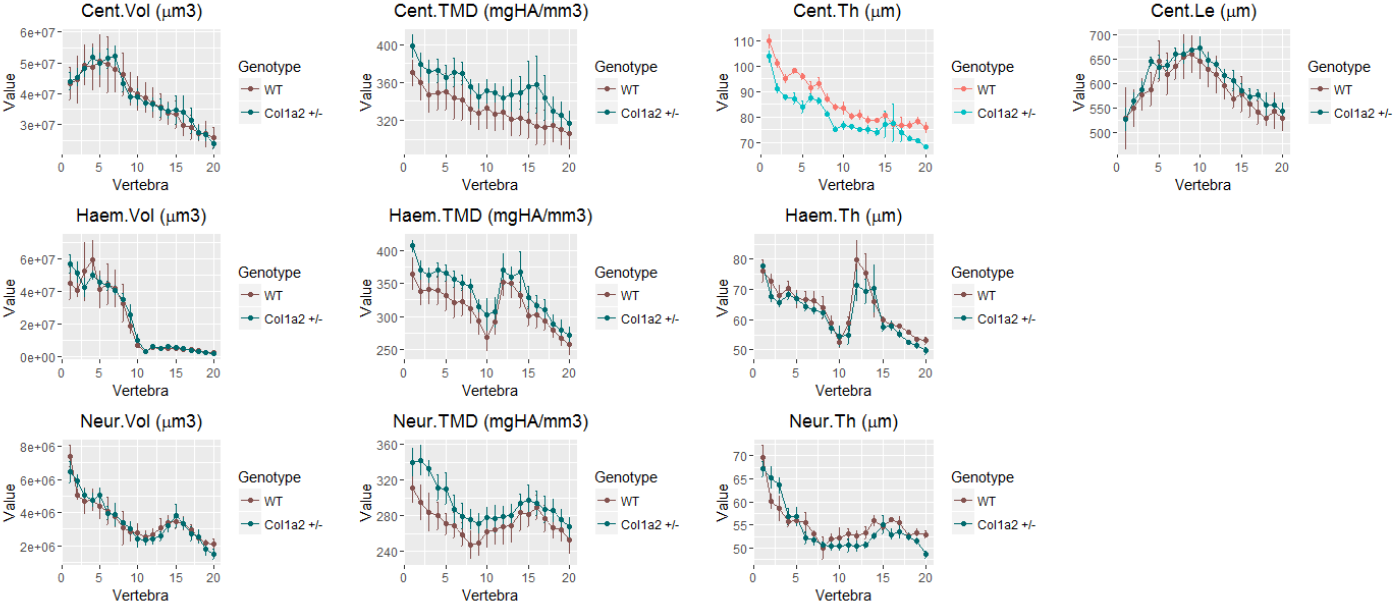
Quantitative µCT-scanning analysis of the vertebral column of *col1a2*^*+/−*^ mutant fish, using FishCuT software. Whole-body µCT scans were acquired in *col1a2* mutant fish and wild type control siblings. FishCuT analysis was performed and interpreted as explained previously (Supplementary Figure 2 legend).

**Supplementary Figure 6.**
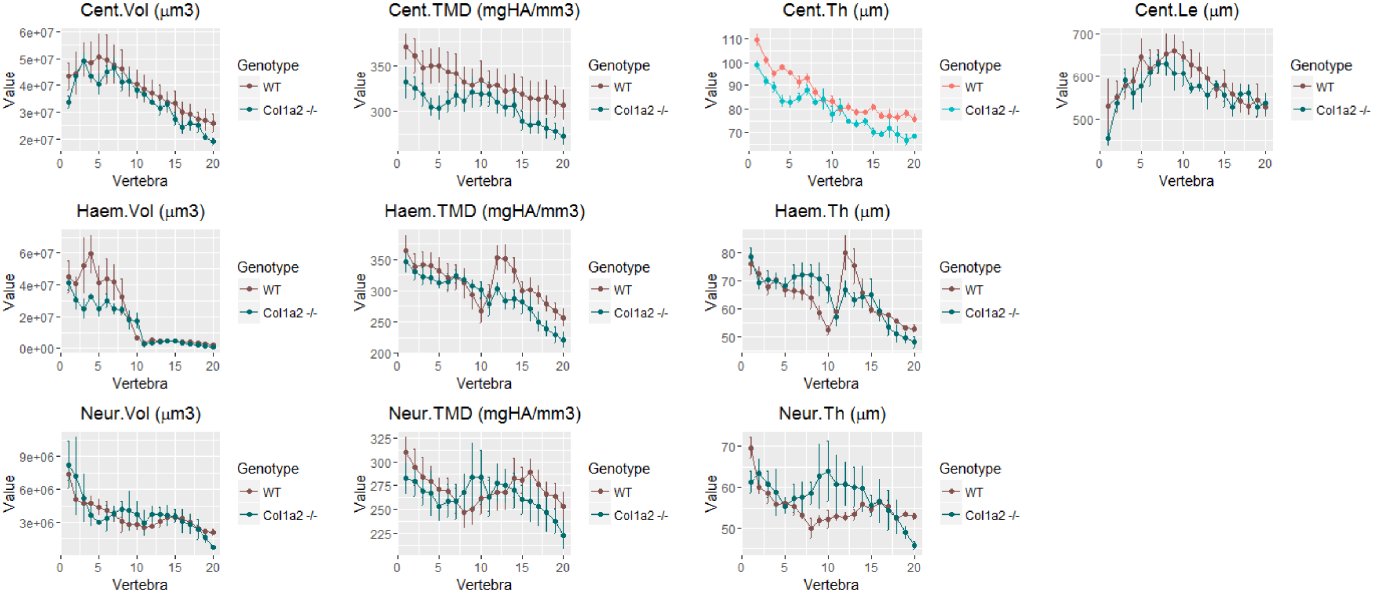
Quantitative µCT-scanning analysis of the vertebral column of *col1a2*^*−/−*^ mutant fish, using FishCuT software. Whole-body µCT scans were acquired in *col1a2*^*−/−*^ mutant fish and wild type control siblings. FishCuT analysis was performed and interpreted as explained previously (Supplementary Figure 1 legend).

**Supplementary Figure 7.**
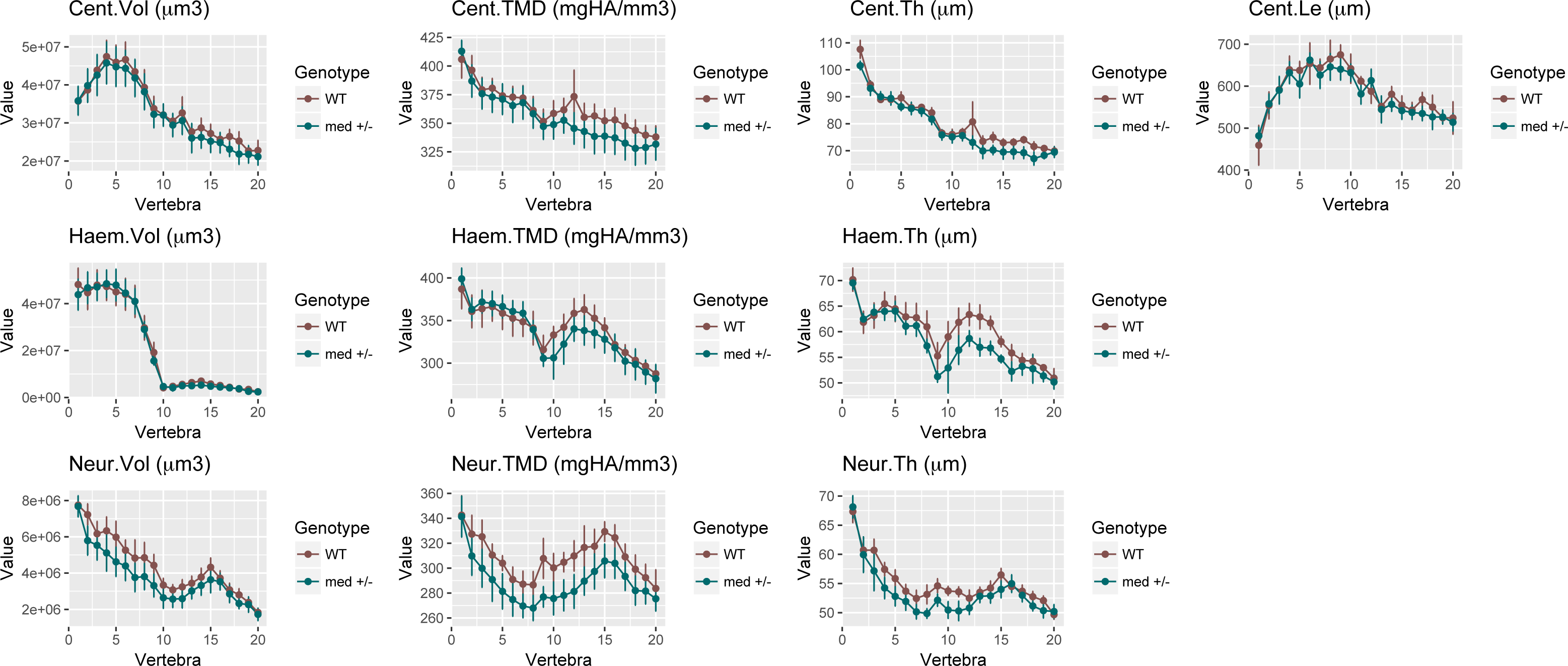
Quantitative µCT-scanning analysis of the vertebral column of *col1a1a*^*med/+*^ mutant fish, using FishCuT software. Whole-body µCT scans were acquired in *col1a1a*^*med/+*^ mutant fish and wild type control siblings. FishCuT analysis was performed and interpreted as explained previously (Supplementary Figure 1 legend).

**Supplementary Figure 8.**
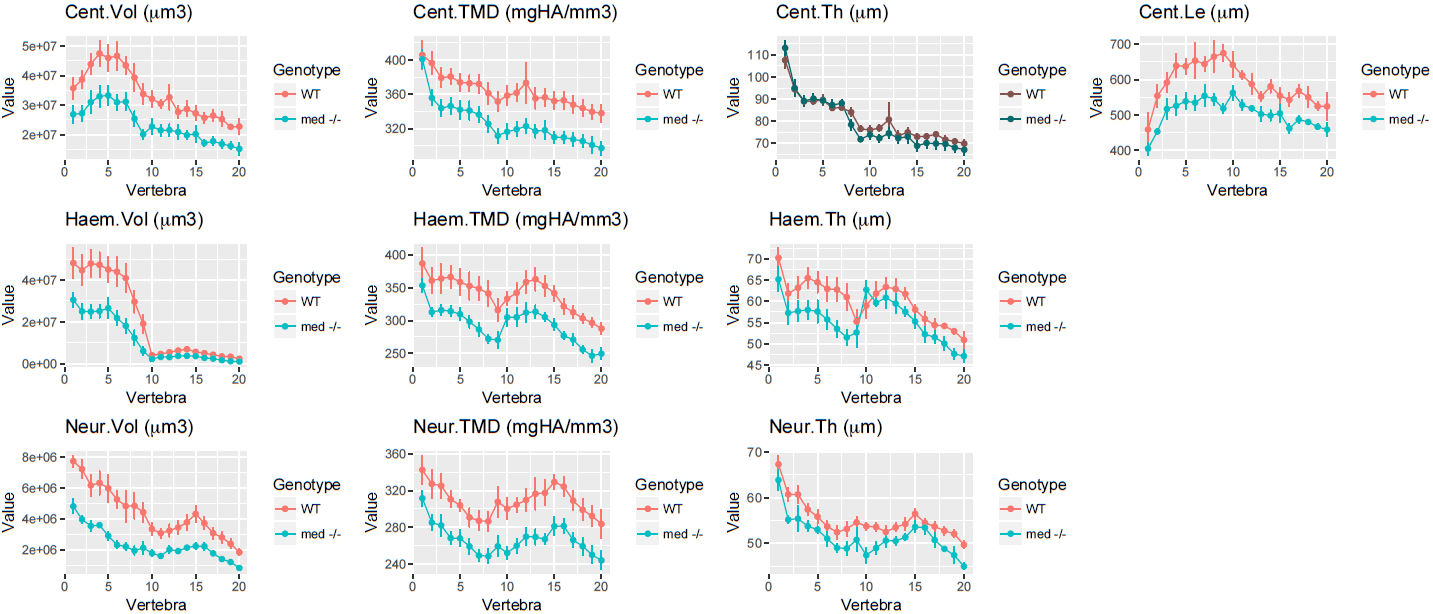
Quantitative µCT-scanning analysis of the vertebral column of *col1a1a*^*med/med*^ mutant fish, using FishCuT software. Whole-body µCT scans were acquired of *col1a1b*^*dhm29/+*^ of *col1a2*^*dhm15/+*^ of *col1a1a*^*dhm14/+*^ of *col1a1a*^*dhm13/+*^ in *col1a1a*^*med/med*^ mutant fish and wild type control siblings. FishCuT analysis was performed and interpreted as explained previously (Supplementary Figure 1 legend).

**Supplementary Figure 9.**
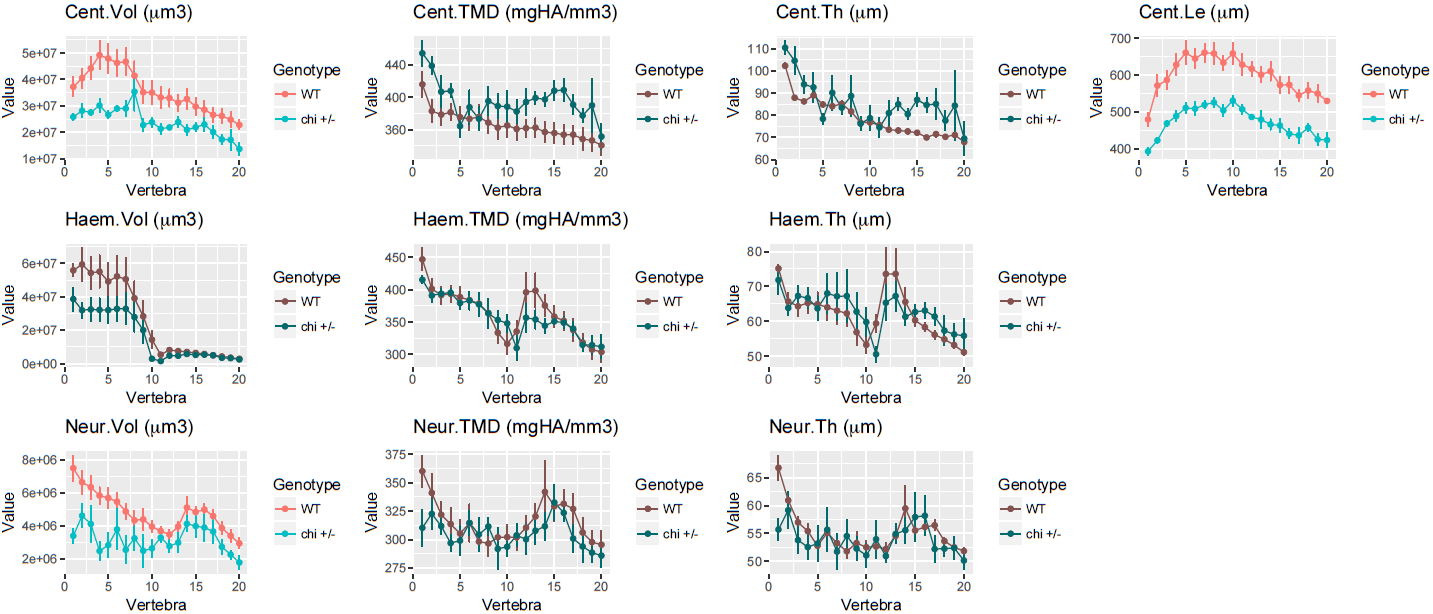
Quantitative µCT-scanning analysis of the vertebral column of *col1a1a*^*chi/+*^ mutant fish, using FishCuT software. Whole-body µCT scans were acquired in *col1a1a*^*chi/+*^ mutant fish and wild type control siblings. FishCuT analysis was performed and interpreted as explained previously (Supplementary Figure 1 legend).

**Supplementary Figure 10.**
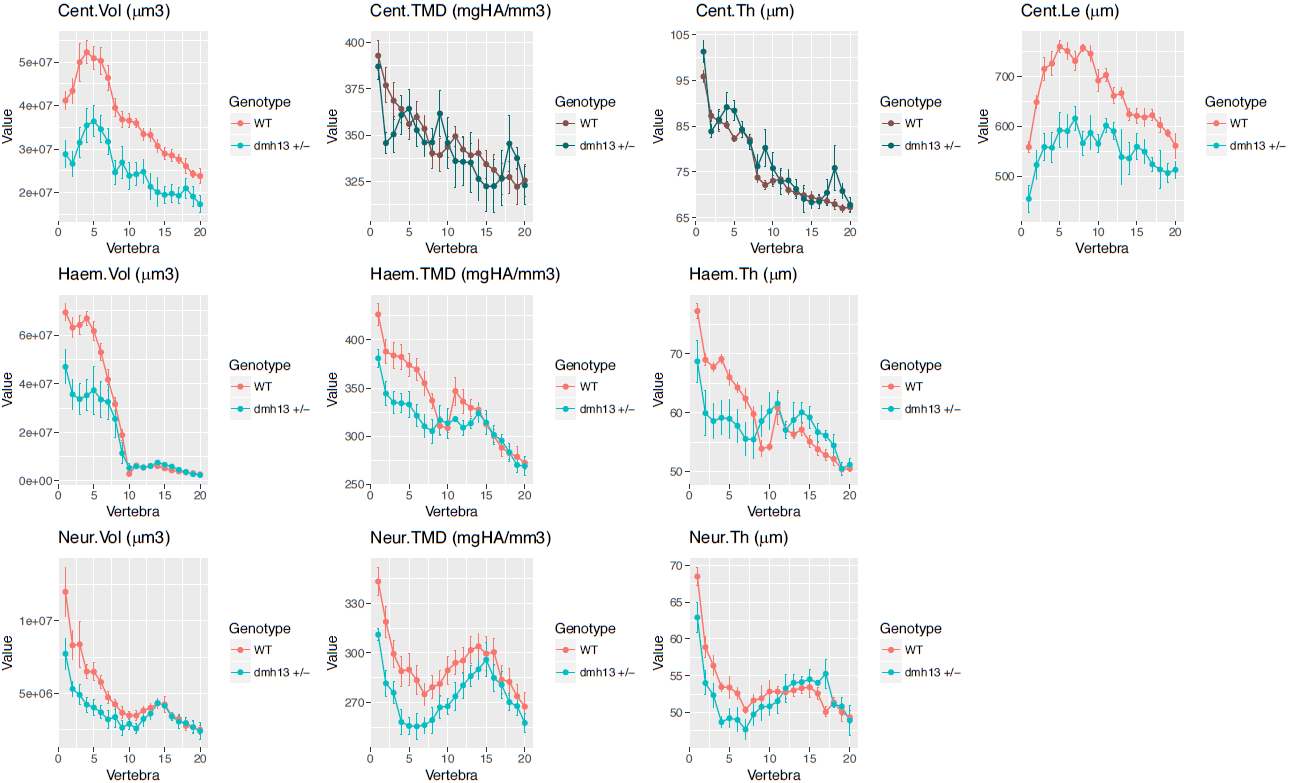
Quantitative µCT-scanning analysis of the vertebral column of *col1a1a^dhm13/+^* mutant fish, using FishCuT software. Whole-body µCT scans were acquired in *col1a1a*^*dhm13/+*^mutant fish and wild type control siblings. FishCuT analysis was performed and interpreted as explained previously (Supplementary Figure 1 legend).

**Supplementary Figure 11.**
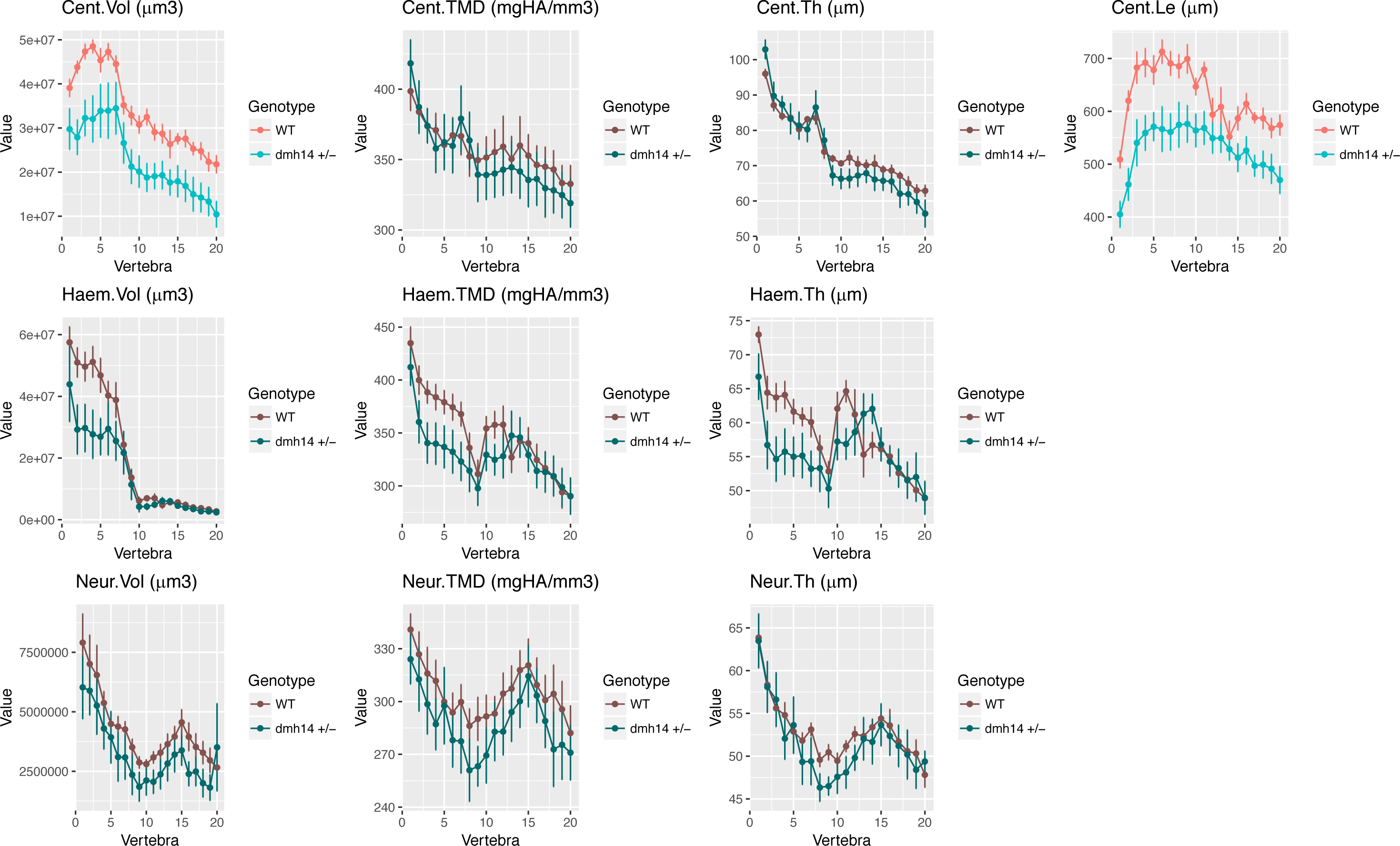
Quantitative µCT-scanning analysis of the vertebral column of *col1a1a^dhm14/+^* mutant fish, using FishCuT software. Whole-body µCT scans were acquired in *col1a1a*^*dhm14/+*^mutant fish and wild type control siblings. FishCuT analysis was performed and interpreted as explained previously (Supplementary Figure 1 legend).

**Supplementary Figure 12.**
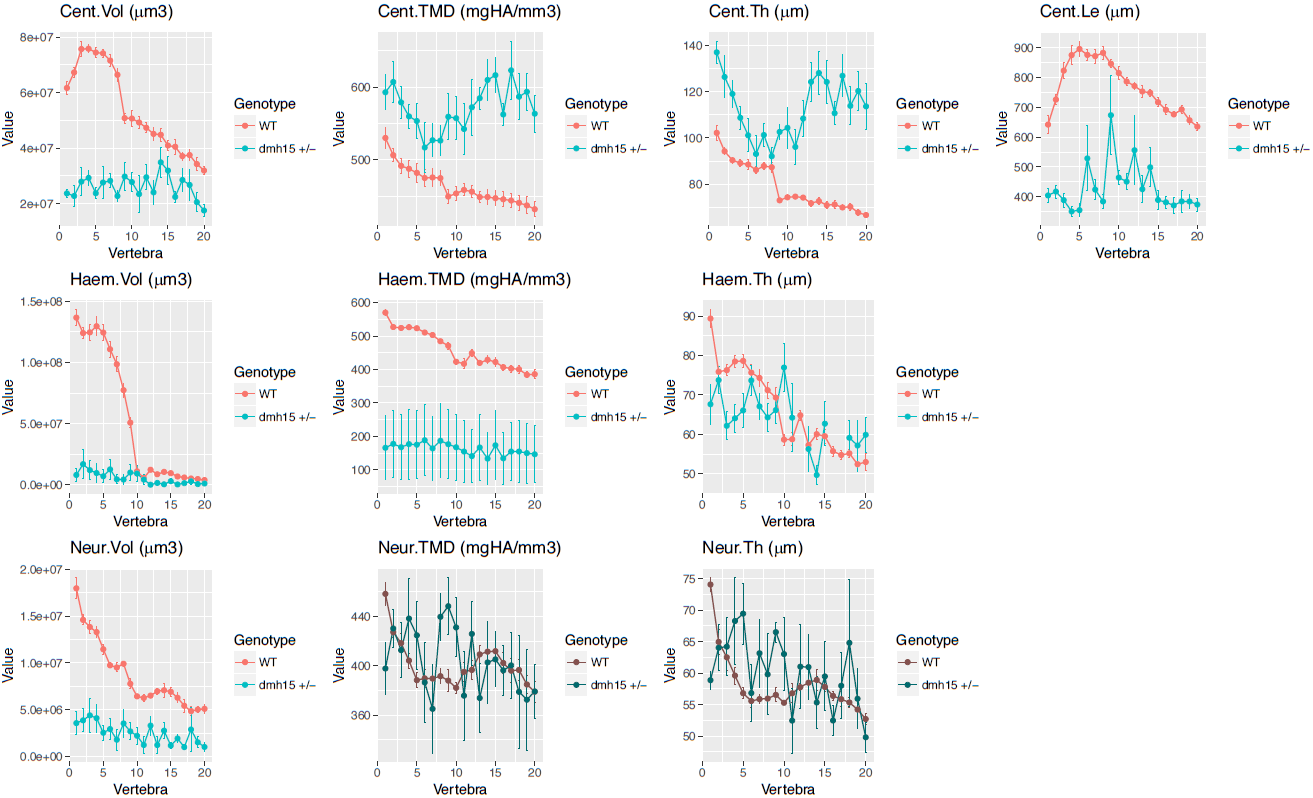
Quantitative µCT-scanning analysis of the vertebral column of *col1a2^dhm15/+^* mutant fish, using FishCuT software. Whole-body µCT scans were acquired in *col1a2*^*dhm15/+*^mutant fish and wild type control siblings. FishCuT analysis was performed and interpreted as explained previously (Supplementary Figure 1 legend).

**Supplementary Figure 13.**
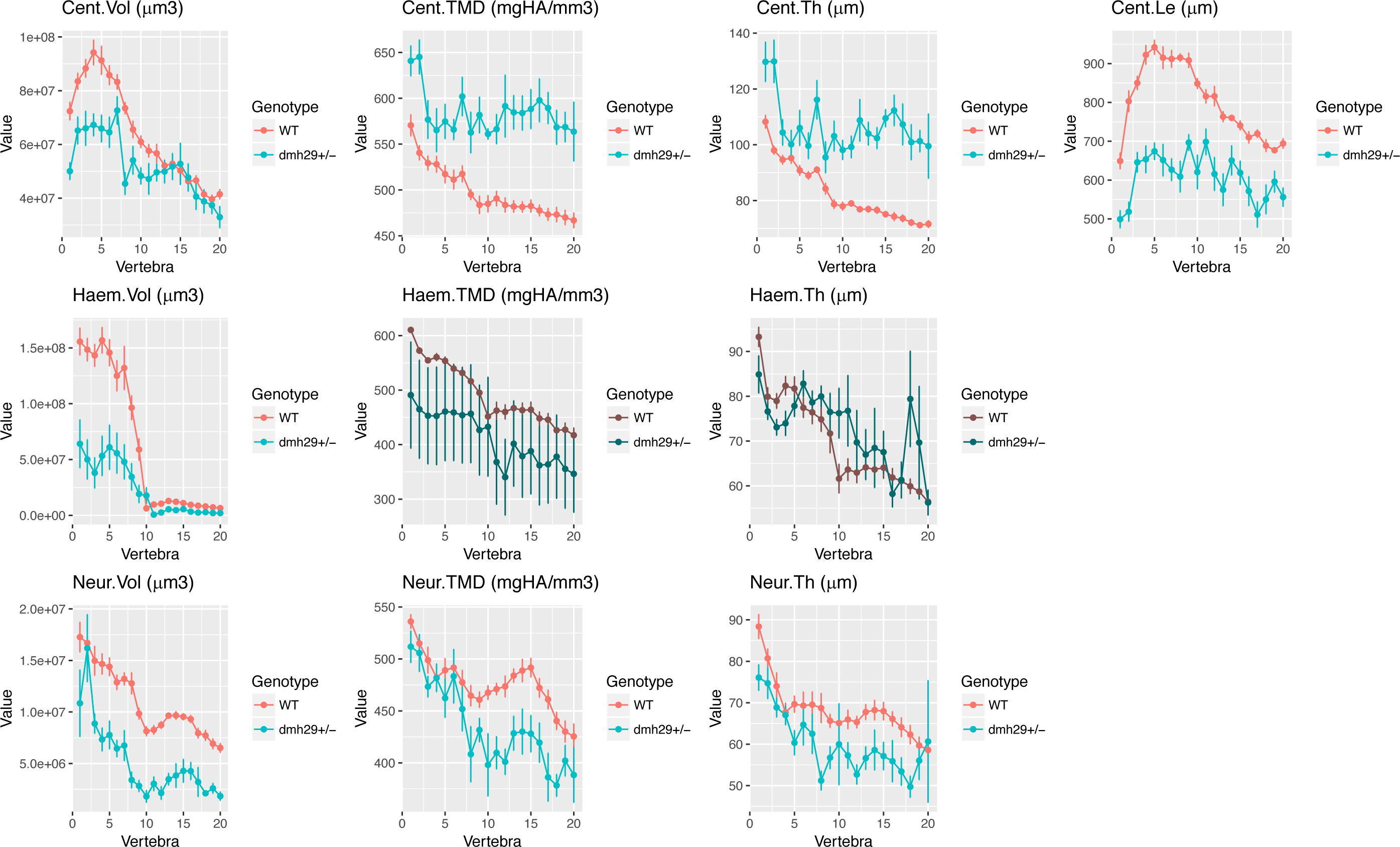
Quantitative µCT-scanning analysis of the vertebral column of *col1a1b^dhm29/+^* mutant fish, using FishCuT software. Whole-body µCT scans were acquired in *col1a1b*^*dhm29/+*^mutant fish and wild type control siblings. FishCuT analysis was performed and interpreted as explained previously (Supplementary Figure 1 legend).

**Supplementary Figure 14.**
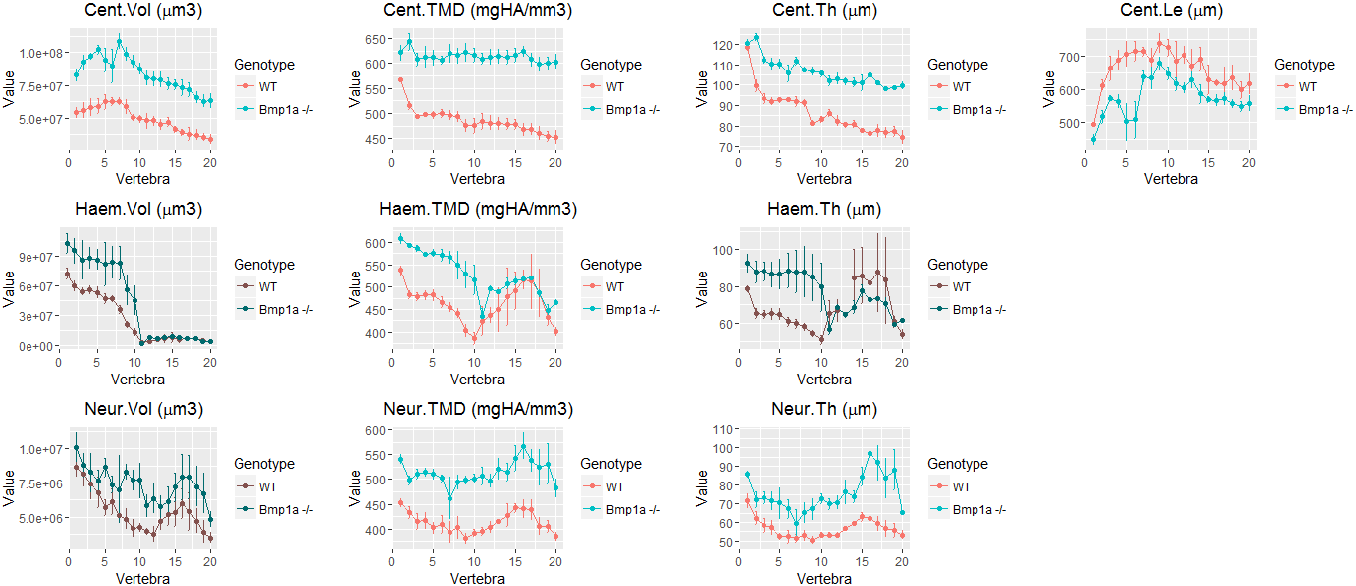
Quantitative µCT-scanning analysis of the vertebral column of *bmp1a*^*−/−*^mutant fish, using FishCuT software. Whole-body µCT scans were acquired in *bmp1a*^*−/−*^mutant fish and wild type control siblings. FishCuT analysis was performed and interpreted as explained previously (Supplementary Figure 1 legend).

**Supplementary Figure 15.**
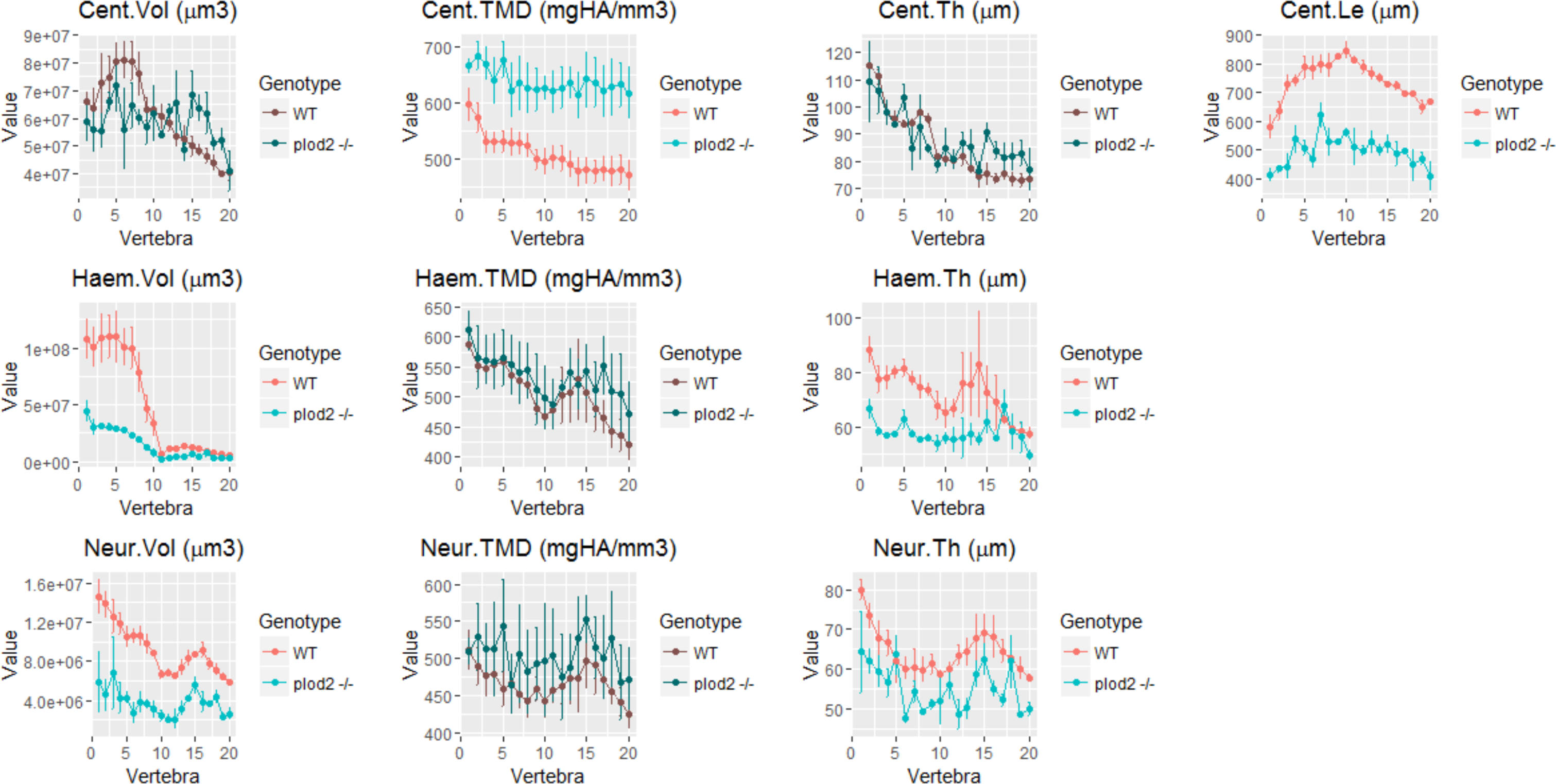
Quantitative µCT-scanning analysis of the vertebral column of *plod2*^*−/−*^ mutant fish, using FishCuT software. Whole-body µCT scans were acquired in *plod2*^*−/−*^ mutant fish and wild type control siblings. FishCuT analysis was performed and interpreted as explained previously (Supplementary Figure 1 legend).

**Supplementary Figure 16.**
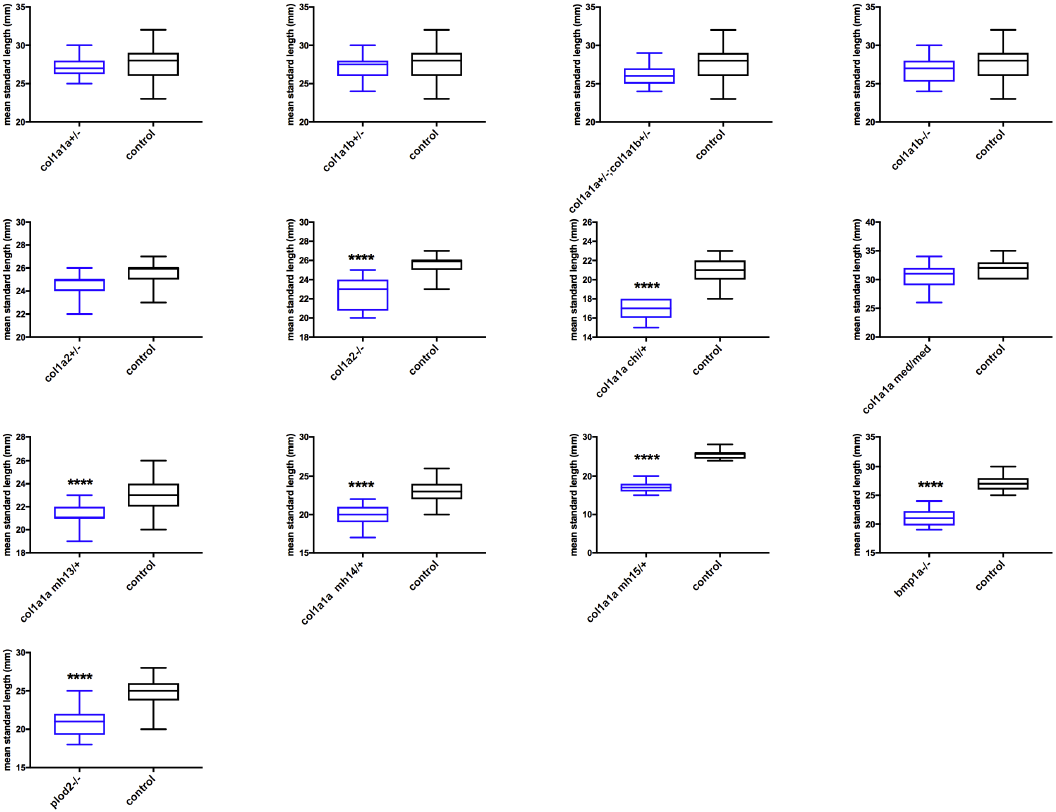
Boxplots of mean standard length per adult mutant genotype. Standard length of adult fish was measured (n=12-20) at adult stage for each mutant genotype and the corresponding control sibling group. P-values were calculated according to the student T-test and means per mutant genotype are reported as being significantly different with p<0.0001 (****).

